# Convergent extension requires adhesion-dependent biomechanical integration of cell crawling and junction contraction

**DOI:** 10.1101/2021.01.12.426405

**Authors:** Shinuo Weng, Robert J. Huebner, John B. Wallingford

## Abstract

Convergent extension is an evolutionarily conserved collective cell movement that elongates the body axis of almost all animals and is required for the morphogenesis of several organ systems. Decades of study have revealed two distinct mechanisms of cell movement during CE, one based on cell crawling and the other on junction contraction. How these two behaviors collaborate during CE is not understood. Here, using quantitative live cell imaging we show that these two modes act both independently and in concert during CE, but that cell movement is more effective when the two modes are integrated via mechano-reciprocity. Based on these findings, we developed a novel computational model that for the first time treats crawling and contraction independently. This model not only confirmed the biomechanical efficacy of integrating the two modes, but also revealed for the first time how the two modes -and their integration- are influenced by cell adhesion. Finally, we use these new insights to further understand the complex CE phenotype resulting from loss of the C-cadherin interacting catenin Arvcf. These data are significant for providing new biomechanical and cell biological insights into a fundamental morphogenetic process that is implicated in human neural tube defects and skeletal dysplasias.

## Introduction

Convergent extension (CE) is a fundamental collective cell movement in which a developing tissue converges along one axis, thereby extending in the orthogonal direction (**Fig. 1A**). CE plays a crucial role in embryogenesis by shaping the body axis during gastrulation and neurulation and by elongating tubular organs during organogenesis (Keller, 2002; Lienkamp et al., 2012; Tada and Heisenberg, 2012). The cell movements of CE are evolutionarily conserved in animals ranging from nematodes and arthropods to vertebrates (Huebner and Wallingford, 2018; Walck-Shannon and Hardin, 2014). Moreover, failure of CE is associated with severe birth defects, including neural tube defects, heart defects, and skeletal dysplasias (Butler and Wallingford, 2017; Wallingford et al., 2013).

**Figure 1.**
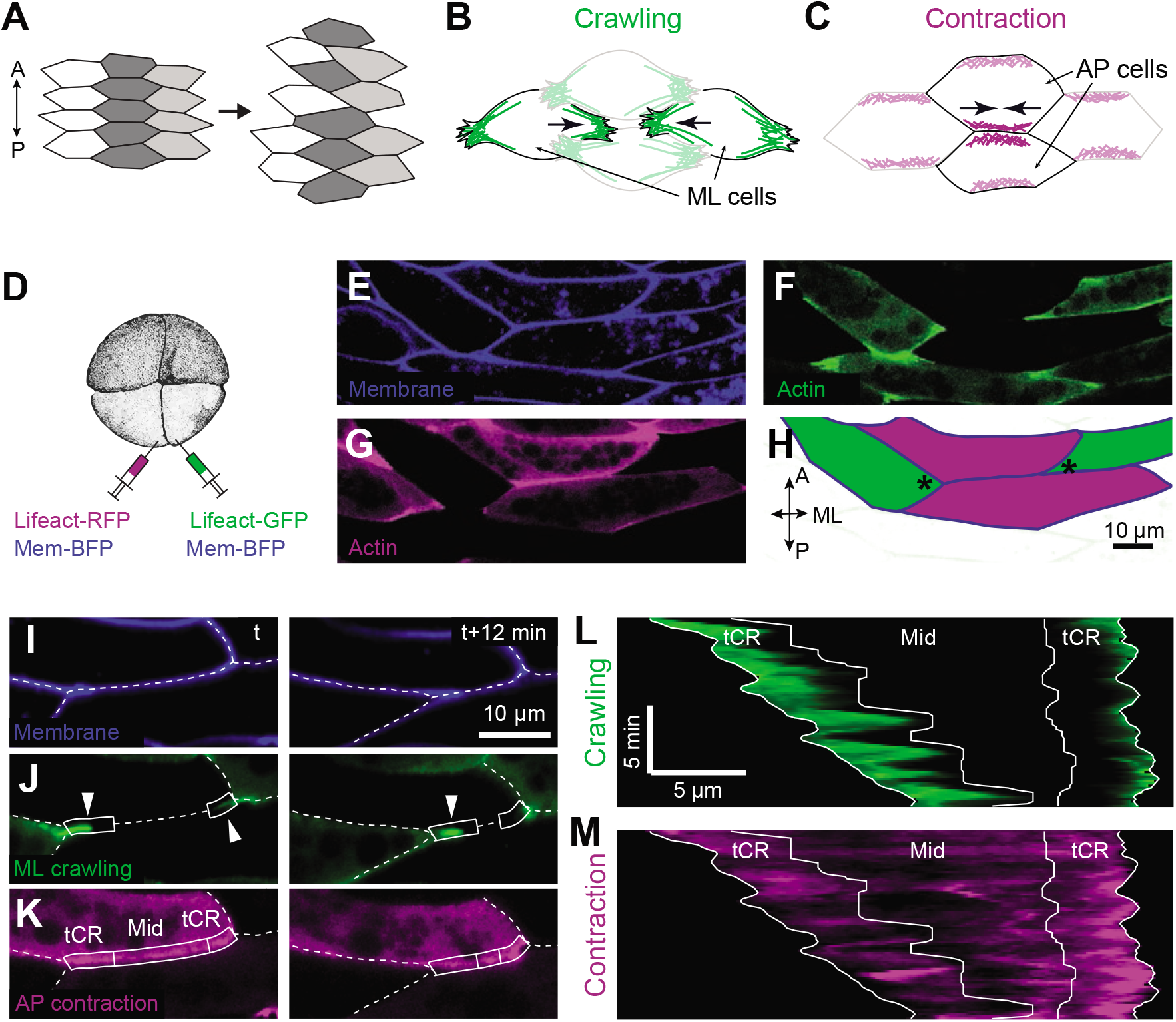
Mosaic labeling showing dynamics of distinct actin populations for crawling and contraction during convergent extension. (A) Illustration of convergent extension showing tissue elongation in the AP axis by cell intercalation in the orthogonal direction. (B) Sketch showing crawling mode of convergent extension in four cells with actin in ML protrusions in green. (C) Sketch showing contraction mode of convergent extension in four cells with actin at AP interfaces in purple. (D) Schematic illustrating mosaic labeling technique in a *Xenopus* embryo. (E-F) Representative images showing uniform labeling of membrane-BFP (E) and mosaic labeling of different colors of an actin biosensor (Lifeact-GFP (F) and Lifeact-RFP (G)). (H) Example showing actin in AP cells labeled in one color and ML cells in another. Asterisks mark representative tCRs for later analysis. (I-K) Still images from a representative time-lapse movie (**Movie S1**) showing membrane (I, blue), actin in the ML protrusions for crawling (J, green, arrowheads) or actin at the AP interface for contraction (K, purple). Dashed lines mark cell-cell interfaces; boxes mark tricellular regions (“tCR”) and the middle region (“Mid”), see text for details. (L, M) Kymograph along the AP cell interface showing spatiotemporal dynamics of actin from the ML protrusions representing the “crawling” signal (L) and actin from the AP cells representing the “contraction” signal (M).

Two distinct cellular mechanisms for convergent extension have been described, and they were initially discovered in different cell types. The first was discovered by work on *Xenopus* body axis elongation during gastrulation (Keller and Hardin, 1987; Keller and Tibbetts, 1989; Shih and Keller, 1992). In this case, intercalation of mesenchymal cells is driven by polarized actin-based protrusions extending from the opposing vertices of mediolateral cells (referred to hereafter as “ML cells”; **Fig. 1B**) (Keller et al., 1992). These protrusions resemble a combination of the sheet-like lamellipodia and spike-like filopodia observed on the leading edge of cultured migrating cells (Devitt et al., 2021), pushing the boundary between neighboring cells and brining ML cells together (**Fig. 1B**). They also form adhesions with the substrate and with the neighboring cells, thus driving cell intercalation in a manner similar to cell migration (Keller and Sutherland, 2020; Pfister et al., 2016).

The second cellular mechanism was discovered in epithelial cells during *Drosophila* germband extension (Irvine and Wieschaus, 1994). In this case, cell intercalation is achieved via polarized junction remodeling, in which junctions joining anteroposterior neighboring cells (referred to hereafter as “AP cells”) shorten to bring together the two dorsoventral neighboring cells.

Actomyosin accumulates on the AP cell interfaces and is activated to provide the contractile force (Bertet et al., 2004; Zallen and Wieschaus, 2004)(**Fig. 1C**). We will refer to these two distinct modes as the “crawling” and “contraction” modes, respectively (**Fig. 1B, C**).

The two modes were initially considered as distinct mechanisms that were implemented in either mesenchyme or epithelia (e.g. (Lienkamp et al., 2012; Nishimura et al., 2012)), cell types that differ significantly in terms of cell-cell adhesion and cell polarity. However, recent evidence suggests that many cell types employ both modes during CE (Huebner and Wallingford, 2018; Shindo, 2018). For example, the two modes were found to work in conjunction in epithelial cells, first in the mouse neural plate, and later in the *Drosophila* germ band (Sun et al., 2017; Williams et al., 2014). In both cases, the crawling mode acts via basolaterally positioned protrusions, while contractions act apically at the epithelial junctions. Interestingly, we previously identified a role for the contraction mode in CE of mesenchymal cells of the *Xenopus* notochord (Shindo et al., 2019; Shindo and Wallingford, 2014), the very cells in which the crawling mode was first defined.

Together, these previous studies suggest that crawling and contraction modes may be integrated in some manner to confer a maximal biomechanical advantage. The nature of such integration is unknown. Here, we used quantitative live-cell microscopy and a novel computational model to demonstrate that cell crawling and junction contraction act both independently and collaboratively to drive CE during *Xenopus* gastrulation. Furthermore, a novel modeling approach suggests that fine control of cell adhesion is essential for mechanoreciprocal integration of crawling and contraction, and experimental manipulation of the cadherin-interacting catenin Arvcf *in vivo* validated this prediction. These data provide new biomechanical and cell biological insights into a fundamental morphogenetic process implicated in human diseases and have broader impacts on studies of collective cell movement in various contexts.

## Results

### Crawling and contraction modes of intercalation act both independently and in concert during vertebrate convergent extension

The mesenchymal cells of *Xenopus* gastrula mesoderm are a key paradigm for studies of CE (Chu et al., 2020; Keller et al., 2003), and while both crawling- and contraction-based cell intercalation mechanisms have been explored, how the two are coordinated remains unclear (Huebner and Wallingford, 2018; Shindo, 2018). We therefore sought to simultaneously assess the contributions of crawling and contraction using explants of the *Xenopus* dorsal mesodermal tissue (“Keller” explants).

We therefore used uniform labeling with membrane-BFP to segment cells and to assess cell intercalation and used mosaic labeling for different colors of an actin biosensor (Lifeact-RFP/Lifeact-GFP) to differentiate populations of actin in neighboring cells (**Fig. 1D-H**; **Movie S1**; **Movie S2**). In this assay, we could clearly distinguish actin dynamics in ML cell protrusions from cortical actin dynamics in adjacent AP cells (**Fig. 1I-K**; **Supp. Fig. 1A-C**). When observed in kymographs, both the ML- and AP-associated actin dynamics were pulsatile and highly heterogeneous, as expected (**Fig. 1L, M**; **Supp. Fig. 1D, E**)(Kim and Davidson, 2011; Pfister et al., 2016; Shindo et al., 2019; Shindo and Wallingford, 2014). These kymographs further suggested that both ML and AP actin dynamics were concentrated in regions near tricellular vertices (**Fig. 1L, M**, “tCR”), while AP pulses outside these regions appeared less frequent and less pronounced (**Fig. 1M**, “Mid”; See Methods for detailed definition of tCR and Mid regions).

We then quantified the contributions of crawling and contraction to cell intercalation. Details are presented in the Methods and in **Supplemental Figures 2, 3**, but briefly: 1) We quantified cell intercalation by measuring the displacement of tricellular vertices connecting AP and ML cells (**Fig. 1I**). 2) We quantified crawling using ML actin intensity as a proxy (**Fig. 1J**). 3) We quantified contraction using AP actin intensity as a proxy (Shindo et al., 2019; Shindo and Wallingford, 2014), treating AP actin dynamics in the tCR and Mid regions independently (**Fig. 1K**). 4) For both displacement and actin dynamics, we quantified each tricellular region separately, as these are known to behave independently in diverse models of intercalation (**Supp. Fig. 4A**)(Cavanaugh et al., 2021; Huebner et al., 2021a; Vanderleest et al., 2018). In bulk analysis, intercalation correlated little with crawling, tCR contraction, or mid contraction (**Supp. Fig. 4B-D**). By contrast, we observed a much stronger correlation between crawling and tCR contraction at each tCR (**Supp. Fig. 4E**), suggesting the possibility of cooperative action.

We considered that crawling and contraction mechanisms could either “take turns” or work together, or both. To explore these possibilities, we took advantage of the pulsatile nature of both cell movement and actin dynamics during CE (**Movie S3**). We first used individual peaks in intercalation velocity curves over time to identify single intercalation “steps” (**Fig. 2A**, left). Then, for each step, we searched for cross-correlated peaks in ML and AP actin intensity within the 40-sec window preceding the velocity peak (**Fig. 2A**, right). This method allowed us to unambiguously associate each step of cell intercalation movement with crawling- and/or contraction-related actin dynamics (e.g. **Supp. Fig. 5A**).

**Figure 2.**
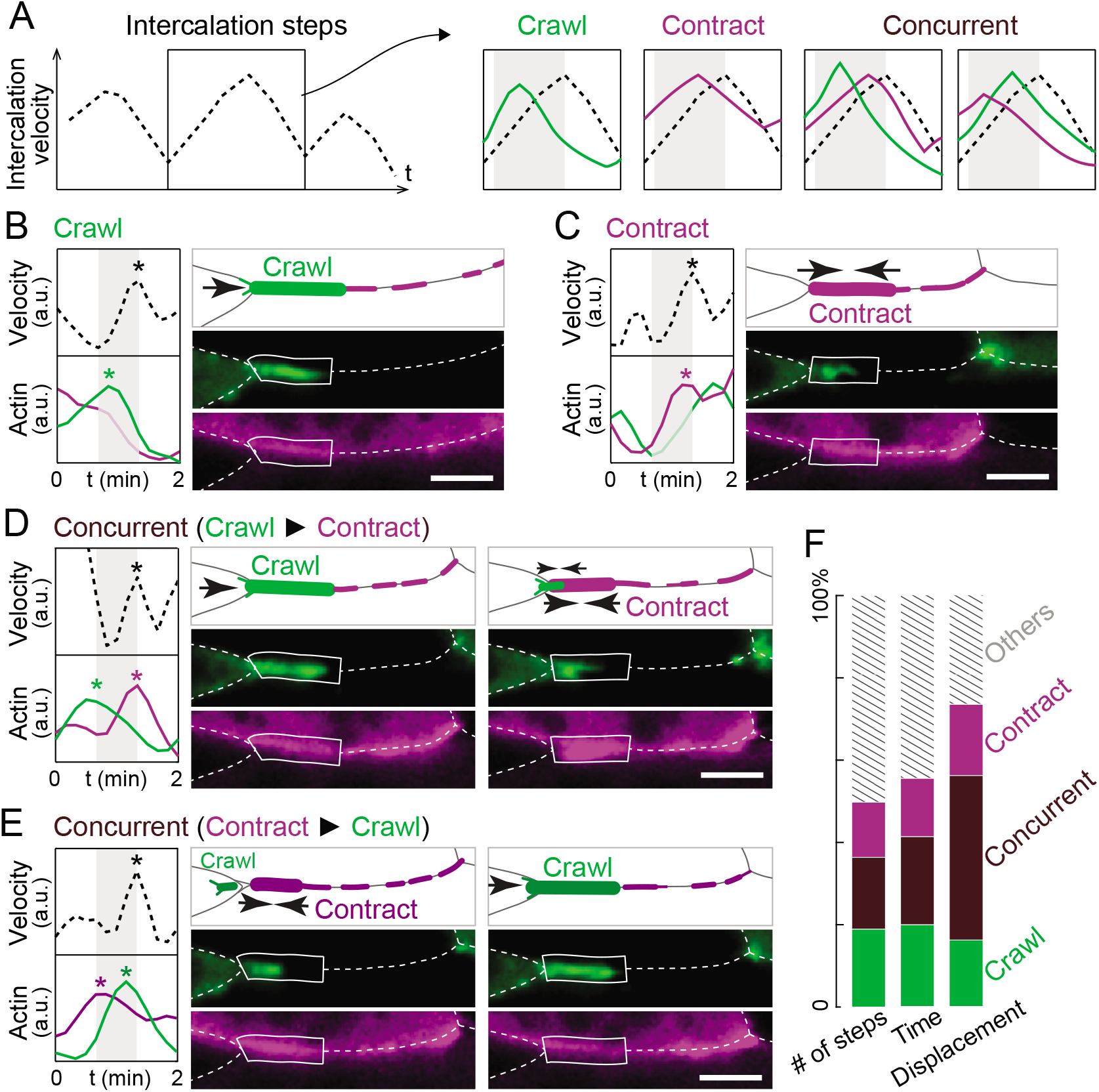
Direct quantification of crawling- and contraction-based cell intercalation during convergent extension. (A) Method for intercalation step analysis. Left, Intercalation steps were identified as individual peaks on the trace of intercalation velocity. Right, each step was categorized based on its correlation with peaks in the crawling and/or contraction signals (green and purple, respectively). Gray boxes mark the 40 sec. time window for correlation analysis. (B-E) Examples of crawling only (B), contraction only (C), and concurrent intercalation steps (D, E). Each shows traces of intercalation velocity and actin dynamics (left), a schematic (top-right), and still frames (bottom-right) from time-lapse data. Asterisks mark the correlated peaks of velocity, crawling, and/or contraction. (F) Stack plots showing in percentile the number of steps, accumulative time of each step, and total displacement of cell intercalation for each category as indicated. Intercalation steps having no correlation with crawling or tCR contraction was labeled as “Others”.

In our data, we identified intercalation steps associated exclusively with ML cell crawling (i.e. correlated with a crawling peak in ML actin intensity; **Fig. 2B**; **Movie S4**) as well as steps associated exclusively with contraction (i.e. correlated with a contraction peak in AP actin intensity; **Fig. 2C**; **Movie S5**). In addition, we identified steps associated with peaks in both ML and AP actin intensity (**Fig. 2D, E**; **Movie S6**; **Movie S7**). These “concurrent” intercalation steps included cases in which a crawling peak precedes a contraction peak (**Fig. 2D**; **Movie S6**), and a similar number of cases in which a contraction peak precedes a crawling peak (**Fig. 2E**; **Movie S7**). We note here that we also observed a small number of intercalation steps associated exclusively with non-tCR contraction from the middle portion of an AP interface (**Supp. Fig. 5B**; **Movie S8**) as well as steps for which no associated crawling or contraction peak could be identified (**Supp. Fig. 5C**, “Others”). Since the vast majority of intercalation can be explained by ML crawling or tCR contraction or both (**Supp. Fig. 5C**), steps in these latter two categories were removed from further analysis.

We found that roughly one-third of steps were associated purely with crawling, another third with contraction, and the final third with both concurrently (**Fig. 2F**, left). Intercalation steps are known to be highly heterogeneous, so it was notable that when we considered the time, rather than the number of steps, we again found an equal distribution of crawling, contraction, and concurrent steps (**Fig. 2F**, middle). By contrast, when we considered actual displacement for cell intercalation, we found that concurrent steps accounted for far more than one-third of the total intercalation displacement moved by tricellular vertices (**Fig. 2F**, right). Together, these data suggest that crawling and contraction can be simultaneously integrated to produce more effective cell movement than can either mode acting alone.

### Concurrent crawling and contraction improves the efficacy of cell intercalation

To further define crawling-only, contraction-only, and concurrent intercalation steps, we quantified several additional metrics. We found that the significantly higher displacement for concurrent steps (**Fig. 3A**) was related to an increase both in the duration of steps and the velocity of vertex movement (**Fig. 3B, C**). Moreover, we noted that some steps involved multiple, successively correlated crawling and contraction peaks (**Fig. 3D**), suggesting an iterative integration of crawling and contraction. We reasoned that if concurrent crawling and contraction provided better efficacy, then multiple concurrent crawling and contraction pulses should be more productive. This was indeed the case, as intercalation accompanied by three or more concurrent crawling and contraction peaks exhibited significantly more displacement and a significantly longer duration than did those accompanied by only two concurrent peaks (**Fig. 3E, F**). Thus, concurrent crawling and contraction produces more effective movement by making each intercalation step both longer-lasting and faster than crawling-only or contraction-only steps.

**Figure 3.**
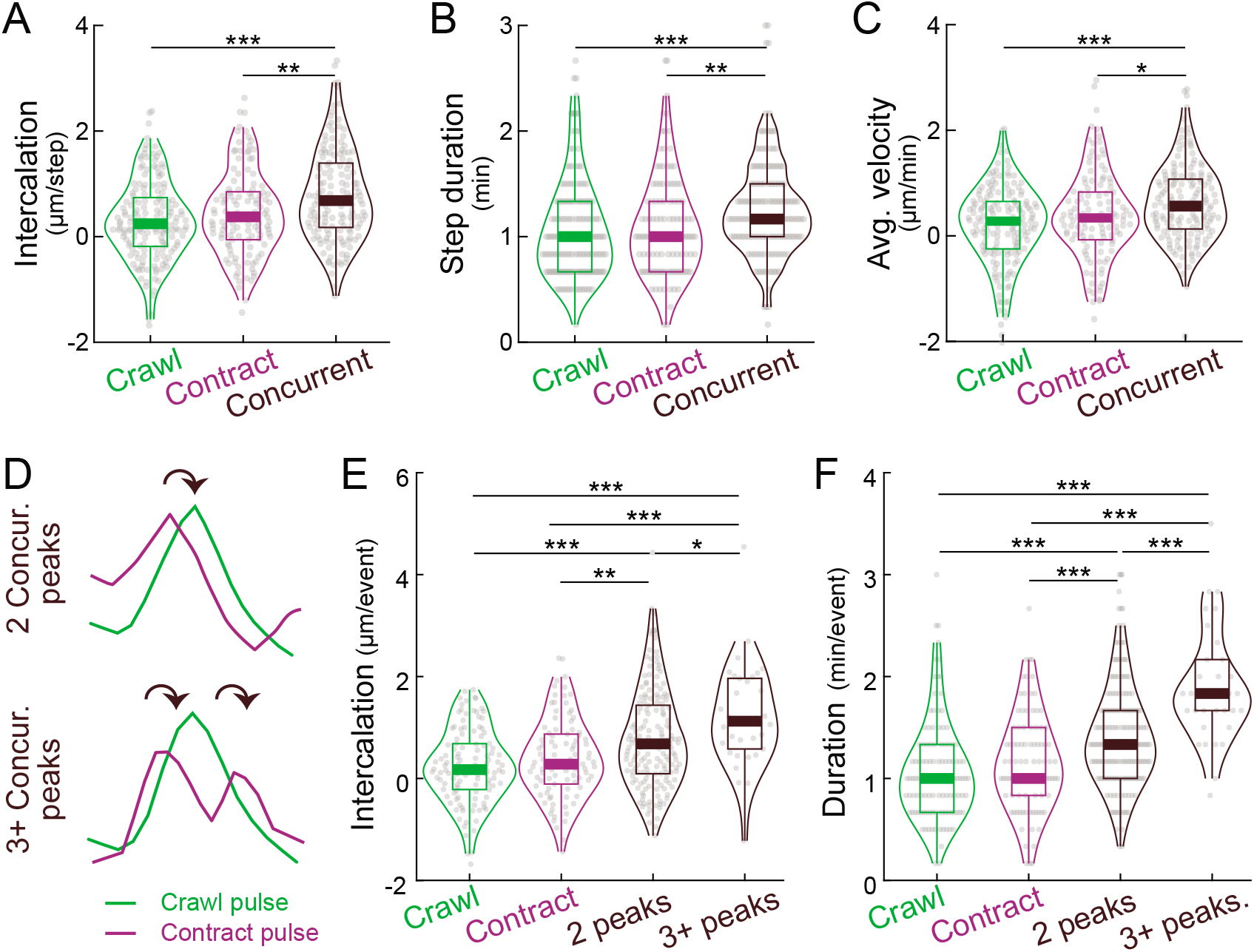
Concurrent crawling and contraction improves the efficacy of cell intercalation. (A) Concurrent steps produce significantly greater intercalation than do crawling or contraction steps. (B, C) Concurrent steps increase both the step duration and average intercalaiton velocity. (D) Sketch showing multiple concurrent crawling and contraction pulses (“3+ Concur. peaks”). (E, F) Multiple concurrent crawling and contraction pulses (“3+ peaks”) further improves the intercalation displacement and duration.

### Concurrent crawling and contraction amplifies actin assembly of both mechanisms

We next sought to understand how concurrent crawling and contraction produced more effective intercalation than does either mode alone. One possibility is biomechanical feedback of actomyosin networks. For example, previous studies in *Drosophila* suggest that tension generated by actomyosin contraction in one cell can stimulate actomyosin contraction in an adherent neighboring cell (e.g. (Fernandez-Gonzalez et al., 2009; Martin et al., 2010). To ask if a similar mechanism was at work here, we first compared contraction-related AP actin dynamics during contraction-only steps with that of AP actin during concurrent steps (**Fig. 4A**). Strikingly, both the peak intensity and the duration of AP actin pulses associated with concurrent steps were significantly amplified compared to contraction-only steps (**Fig. 4B, C**).

**Figure 4.**
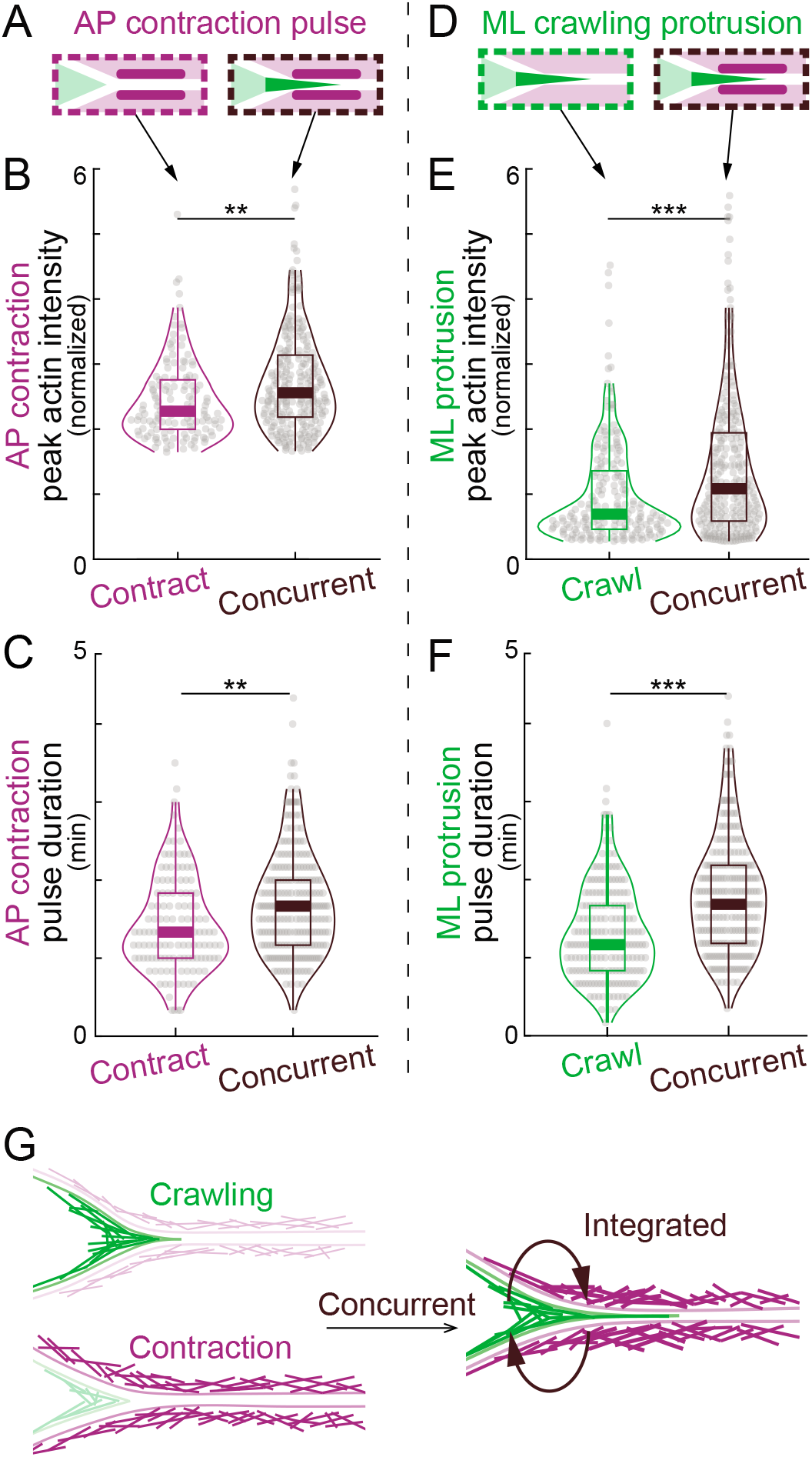
Integration of crawling and contraction enhances actin assembly. (A) Schematic showing contraction pulses (purple) in contraction-only and concurrent steps. (B, C) Violin plots showing increased peak actin intensity (B) and pulse duration (C) of contraction pulses when they occurred concurrently with crawling pulses compared with contraction pulses occurring alone. (D) Schematic showing crawling protrusions (green) in crawling-only and concurrent steps. (E, F) Increased peak actin intensity (E) and pulse duration (F) of crawling protrusions when they occurred concurrently with contraction pulses compared with crawling protrusion occurred alone. (G) Schematic showing amplified actin assembly in crawling protrusions and contraction cortex when they are integrated during concurrent steps.

We performed a complementary analysis of crawling-related ML actin dynamics (**Fig. 4D**), and we observed a similar trend; the peak intensity and duration of crawling protrusions were amplified in concurrent steps relative to crawling-only steps (**Fig. 4E, F**). Thus, concurrent execution of crawling and contraction has a synergistic effect, enhancing actin assembly of both modes. This result suggests that the concurrent steps are, in fact, integrated and that mechano-reciprocity amplifies forces when crawling and contraction occur concurrently (schematized in **Fig. 4G**).

### A novel vertex model recapitulates integrated crawling- and contraction-based convergent extension

Theoretical modeling is a crucial tool in studies of morphogenesis, as it allows manipulation of attributes that may be difficult or impossible to manipulate experimentally, and several modeling studies have been used to explore the mechanisms of convergent extension (e.g. (Alt et al., 2017; Fletcher et al., 2017; Merkel and Manning, 2017)). However, the vertex models commonly employed for such studies are limited, because *a*) they generally treat two sides of a cell-cell interface as a single feature; *b*) heterogeneity along a single cell-cell interface is generally ignored; and *c*) they do not independently consider contributions from cell crawling and junction contraction (Belmonte et al., 2016; Brodland, 2006; Finegan et al., 2019; Shindo et al., 2019).

To overcome these limitations, we re-envisioned the vertex model of convergent extension. We represented each cell not as a six-vertex hexagon, but instead as a 90-vertex polygon, allowing us to model local events in discrete regions of individual cell-cell interfaces (**Fig. 5A**). Furthermore, cell-cell interfaces *in vivo* are formed of two apposed cell membranes, which are linked by cell-cell adhesion molecules but can behave independently. We therefore modeled junctions between two cells as two independent entities (**Fig. 5B**), while modeling cell adhesion to connect adjacent cells (**Fig. 5C**).

**Figure 5.**
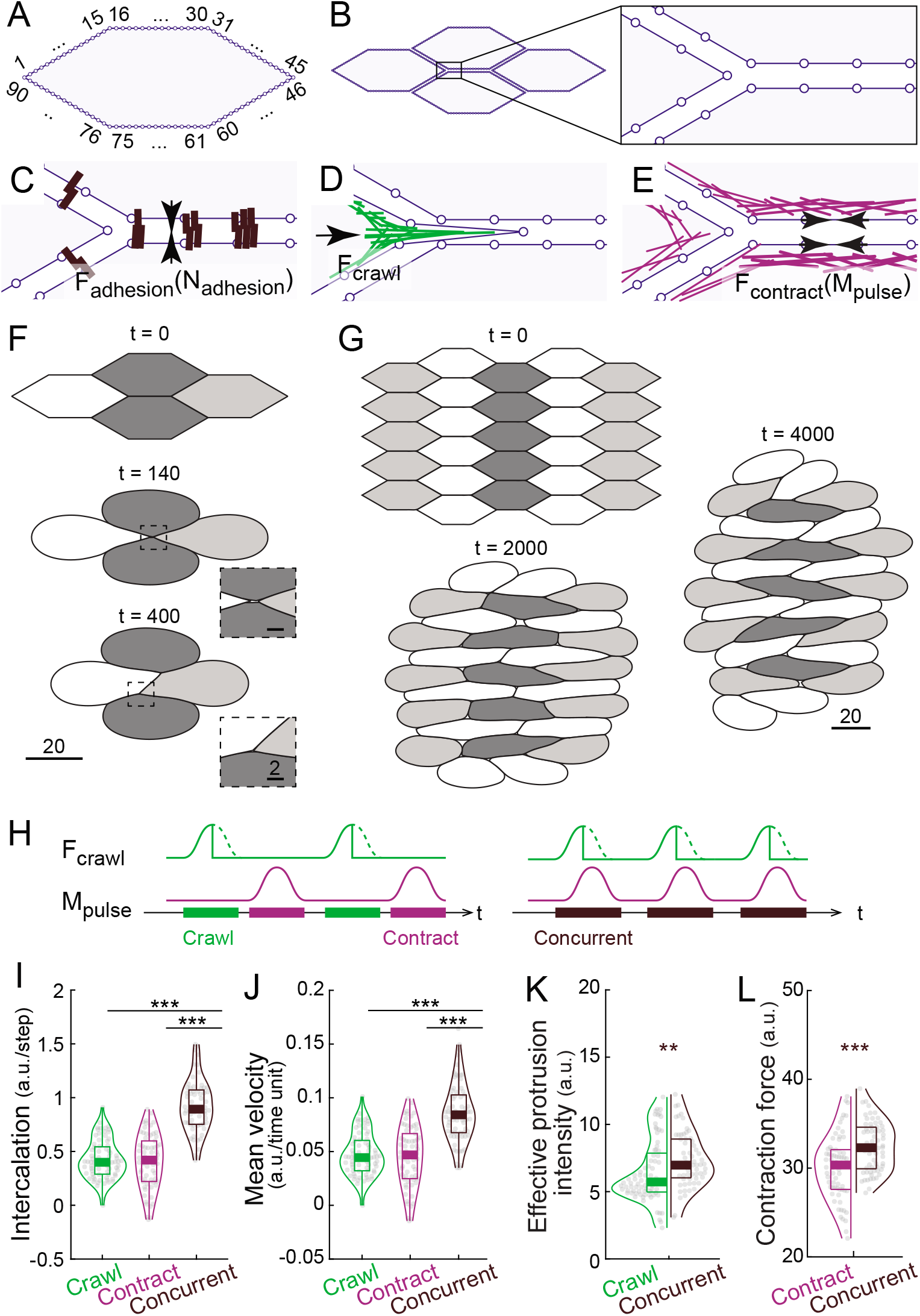
A novel vertex model provides insights into the biomechanics of crawling-contraction integration. (A) Each individual cell was modelled as a 90-vertex polygon. (B) A 4-cell model, with cell-cell interfaces modeled as independent entities. (C-E) Schematic focused on the boxed region in (B), showing the design for subcellular modeling behaviors. (C) Cell-cell interfaces were connected via cell adhesion clusters of *N_adhesion_*, holding adhesive forces of *F_adhesion_*. (D) Crawling forces *F_crawl_* were applied to ML vertices around tricellular regions. (E) Contraction forces *F_contract_* were added to cell cortex, which is a function of the amount of actomyosin density including the pulsatile component *M_pulse_*. (F) Representative simulation result of a 4-cell model; magnification in insets reveals minimal extracellular gaps between cells. (G) Representative simulation result of a 27-cell model, showing not only cell intercalation but also tissue-wide convergent extension. (H) Schematic showing timepoints for crawling forces and contraction pulses, such that crawling-only or contraction-only intercalations steps can be distinguished from concurrent steps. (I, J) The model recapitulates enhanced intercalation displacement and higher velocity for concurrent steps as compared to crawling or contraction alone. (K, L) Violin plots showing synergistic effect of integration on effective protrusion intensity (K) and AP contraction force (L), recapitulating experimental observations (see Fig. 4).

Details of the model are presented in the Methods, but briefly, we invoke three key forces: First, the pushing force for cell crawling was modeled as a defined force profile applied on tricellular vertices (**Fig. 5D**, “*F_crawl_*”). Second, the contractile force from actomyosin pulses (“*M_pulse_*”) for junction contraction was modeled with Hill’s muscle model (Mitrossilis et al., 2009) (**Fig. 5E**, “*F_contract_*”). Finally, we modeled the force transmitted between neighboring cells via force-dependent cell adhesion at cell interfaces using a catch-slip bond model (**Fig. 5C**, “*F_adhesion_*”, “*N_adhesion_*”), which reflects the known role of Cadherin adhesion in convergent extension (Brieher and Gumbiner, 1994; Fagotto et al., 2013; Huebner et al., 2021a; Pfister et al., 2016). Without loss of generality, this model is dimensionless, but parameters for the key force components were estimated from experimental data, with their relative values in a physiologically relevant range (Supp. Table.1).

Critically, our model successfully recapitulated many gross aspects of convergent extension. For example, we first modelled cell intercalation using four-cell models (**Fig. 5F**). To reproduce actin dynamics observed *in vivo* (**Fig. 1L, M**), medially-directed crawling forces (“*F_crawl_*”) and heterogenous actomyosin pulses along the AP cell interface (“*M_pulse_*”) were applied at randomized timepoints (**Supp. Fig. 6A, B**). This configuration consistently produced complete cell intercalation (i.e., “T1 transition”), including shortening of the AP cell interface and formation and elongation of a new AP directed cell-cell interface (**Fig. 5F**; **Movie S9**). We also demonstrated using our model that contraction from the middle portion of an AP cell interface has limited effect on cell intercalation at tCRs (**Supp. Fig. 6D**), consistent with *in-vivo* data (**Supp Fig. 5C**).

Moreover, 27-cell clusters in which both medially and laterally directed crawling forces were applied to each cell at equal probability consistently recapitulated not just cell intercalation, but also tissue-wide convergent extension (**Fig. 5G**; **Movie S10**). Importantly, cells were able to accomplish multiple rounds of cell intercalation and interdigitated in a regular manner (**Fig. 5G**). Future cell elongation in the ML axis with cell intercalation was also observed, similar to the reported cell shape change *in vivo* (Wilson et al., 1989). Thus, a combination of crawling and contraction in our novel model can recapitulate cell movements and tissue morphology change observed in vertebrate convergent extension *in vivo*.

### Modeling recapitulates biomechanical synergy of crawling/contraction integration

As a further test of our model’s validity, we asked if it could also recapitulate finer-scale behaviors observed *in vivo*. To this end, we used four cell models to simulate shortening of AP interfaces. We set medially directed crawling forces and AP actomyosin pulses in tCRs to defined time points so that we could distinguish crawling-only or contraction-only steps from steps with concurrent crawling and contraction (**Fig. 5H**). Importantly, we found that across a wide range of parameters for *F_crawl_* and *M_pulse_*, concurrent crawling and contraction in the model elicited significantly greater displacement and mean intercalation velocity (**Fig. 5I, J**; **Supp Fig. 7A, B**), similar to the effects observed for concurrent steps *in vivo*.

This finding prompted us to ask if our model also recapitulated the amplification of actin assembly during concurrent steps that we observed *in vivo* (**Fig. 4**). We defined effective protrusion intensity in our model as the integral of protrusion length over time and used it as a proxy for protrusion dynamics (**Supp. Fig. 6E**). We reasoned that this is a mechano-responsive measure for a given crawling force, reflecting the interaction between a protrusion and its adjacent cells by considering both protrusion extension and retraction (**Supp. Fig. 6E**). We found that when crawling and contraction occurred concurrently, the effective protrusion intensity was significantly higher than when a crawling force was applied alone (**Fig. 5K**). This effect held for a wide range of crawling forces and actomyosin pulses (**Supp. Fig. 7C, D**). Importantly, this mechano-reciprocity was not explicitly designed in the model, but nonetheless recapitulated that observed *in vivo* (i.e. **Fig. 4D-F**).

Similarly, our model imposed actomyosin pulses (“*M_pulse_*”) for contraction, and the corresponding contraction forces (“*F_contract_*”) can be estimated directly from the Hill’s muscle model (see Method; example in **Supp. Fig. 6B, C**). Mechanical forces are known to promote assembly of actomyosin structures, forming a feedforward loop in several contexts, including CE (Fernandez-Gonzalez et al., 2009; Martin et al., 2010; Miao and Blankenship, 2020; Uyeda et al., 2011; Zaidel-Bar et al., 2015). We therefore used the calculated contraction force as a proxy for actin dynamics at the AP cell cortex. Over a wide range of crawling forces and actomyosin pulses, we observed amplified contraction forces when crawling and contraction occurred concurrently (**Fig. 5L**, **Supp. Fig. 7E, F**). Because this is not an explicit feature in the model, it suggests a mechano-reciprocal integration of crawling and contraction similar that observed *in vivo* (**Fig. 4A-C**).

Thus, our model recapitulated not only the tissue-scale and cell-scale convergent extension, but also the synergy of crawling/contraction integration for both cell intercalation and enhanced actin assembly. The synergistic effects observed both *in vivo* and *in silico* suggest a mechanoreciprocity between the two distinct populations of actomyosin that drives integration of the crawling and contraction modes of CE.

### Modeling provides insight into the role of regulated adhesion in crawling, contraction, and their integration

Our working model for crawling/contraction integration schematized in **Fig. 4G** suggested a strong dependency on cell adhesion for effective integration. Thus, we next examined the effect of cell adhesion by modulating the total adhesion units available, 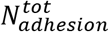, while maintaining a constant, moderate crawling force, *F_crawl_* and constant moderate actomyosin pulses, *M_pulse_*.

Interestingly, when we reduced adhesion by half, we still consistently observed complete cell intercalation in 4-cell models (**Fig. 6A**), though it was characterized by enlarged extracellular voids between cells, particularly in tricellular and quad-cellular regions, reflecting the effect of reduced cell adhesion (**Fig. 6A**, insets). More strikingly, when we simulated convergent extension in a 27-cell model with reduced adhesion, tissue-wide convergent extension was substantially reduced, intercellular voids were enlarged, and cell packing was much less regular (**Fig. 6B**).

**Figure 6.**
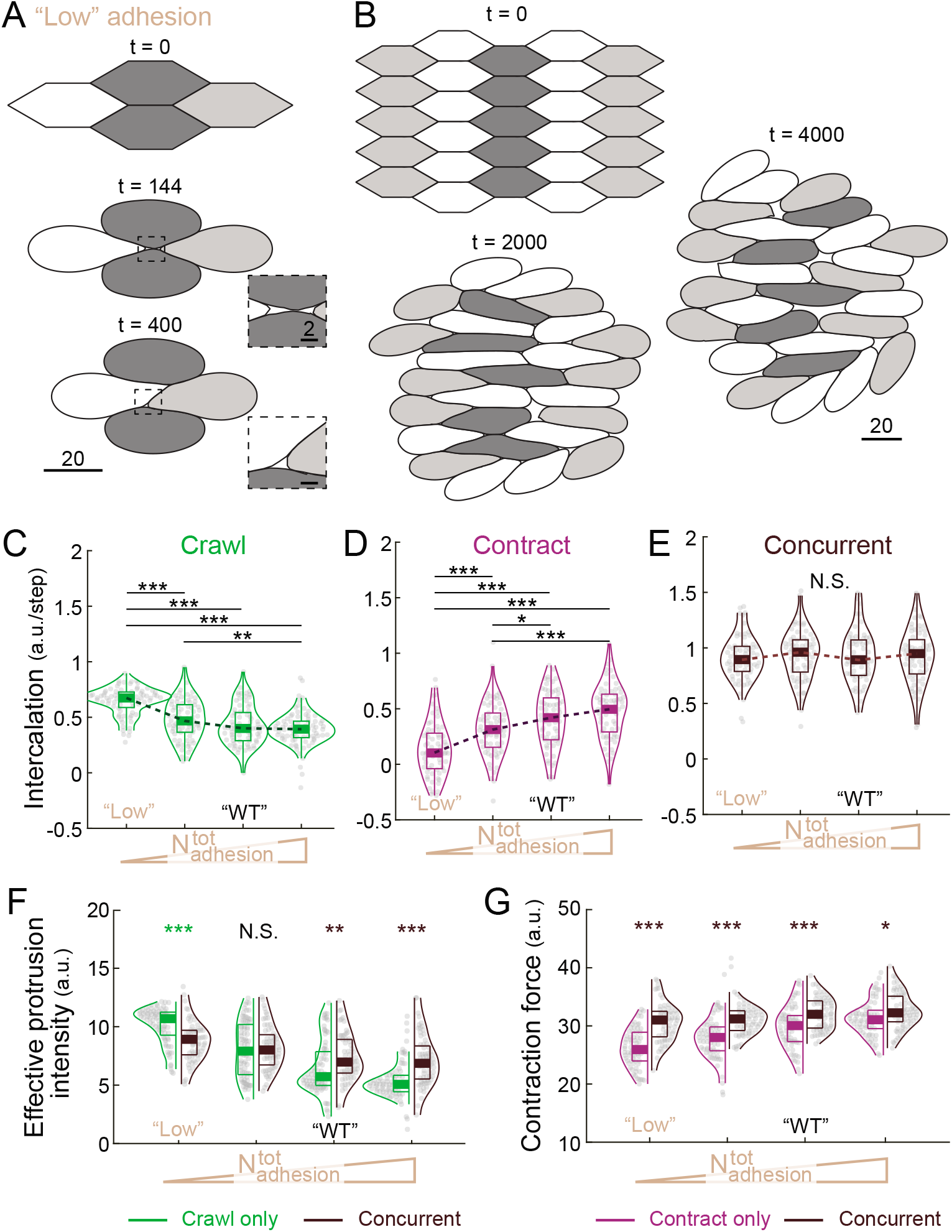
Role of cell adhesion in crawling and contraction. (A) Representative simulation result of a 4-cell model with low adhesion showing complete cell intercalation. Magnification in insets shows enlarged voids in multi-cellular regions. (B) Simulation result of a 27-cell model with low adhesion reduced tissue-wide convergent extension. (C-E) Effect of increasing adhesion on intercalation (vertex displacement) during crawling only, contraction only, and concurrent steps in the model. “WT” marks the value used for simulating the wildtype condition in Fig. 5. “Low” marks the value used for simulating the low-adhesion condition in Fig. 6A, B. (F) Effect of increasing adhesion on the effective protrusion intensity in crawling only and concurrent steps. Enhanced protrusion intensity during concurrent steps is only observed with medium-high adhesion. (G) Effect of adhesion on contraction force in contraction only and concurrent steps. Contraction force during concurrent steps is robustly increased in concurrent steps.

We then used the modeling approaches outlined above (i.e., **Fig. 5H**) to ask how modulation of adhesion impacts on different modes of cell intercalation. We found that higher adhesion strongly inhibited crawling-only intercalation while favoring contraction-only intercalation (**Fig. 6C, D**, “Low” vs. “WT”). It is also notable that these effects plateaued when adhesion strength increased further. Surprisingly, across a wide range of adhesion strength, elevated intercalation with concurrent crawling and contraction was maintained robustly (**Fig. 6E**).

We next examined the impact of adhesion on integrating concurrent crawling and contraction. We found that effective protrusion intensity tended to decrease as adhesion increased (**Fig. 6F**), a result similar to cadherin mediated contact inhibition of locomotion *in vivo* (Becker et al., 2014). More interestingly, amplification of protrusion intensity during concurrent steps was only observed when adhesion was in the medium-high regimes (**Fig. 6F**). In low-adhesion regimes, protrusion was indeed diminished in concurrent steps as compared to crawling-only steps (**Fig. 6F**, “Low”), reflecting the requirement for adhesion between protrusions and the cortex of another cell to interact. In contrast, there was a pronounced trend for the contraction force to increase with adhesion (**Fig. 6G**), consistent with the thought that anchoring sites are necessary for the actomyosin to generate cortical tension. Interestingly, the synergistic effect on contraction force during concurrent steps was quite robust to changes in adhesion (**Fig. 6G**), suggesting a differential impact of adhesion and the transmitted force on contraction versus crawling.

### Multiple functions of the Arvcf catenin are required for normal integration of crawling and contraction

Finally, we considered that this new quantitative understanding of the interplay of crawling and contraction during of CE might shed light on complex experimental loss-of-function phenotypes. For example, the Arvcf catenin is a C-cadherin/Cdh3-interacting protein that is required for *Xenopus* CE (Fang et al., 2004; Paulson et al., 2000). In a companion paper, we show that loss of Arvcf decreased the tissue-level extension force by cell intercalation, thus leading to convergent extension defect in embryos (Huebner et al., 2021b). The cell biological basis of this phenotype is not known, so we turned to our mosaic labelling approach and our new model for insight.

Live imaging of mosaically labeled explants revealed that Arvcf loss (see methods) did not affect cell shape (**Fig. 7A**) but did decrease overall intercalation velocity (**Fig. 7B**). Interestingly, analysis of individual intercalation steps revealed that crawling-only and contraction-only steps were not affected (**Fig. 7C**), but in striking contrast, intercalation during concurrent steps was significantly reduced in the absence of Arvcf (**Fig. 7C**, right). This mosaic analysis further revealed that the defect in concurrent steps was associated specifically with a failure to amplify crawling-associated actin dynamics during concurrent steps contraction-associated dynamics remained significantly amplified (**Supp. Fig. 8A-D**). This result suggested a specific role for Arvcf in the normal synergistic integration of crawling and contraction.

**Figure 7.**
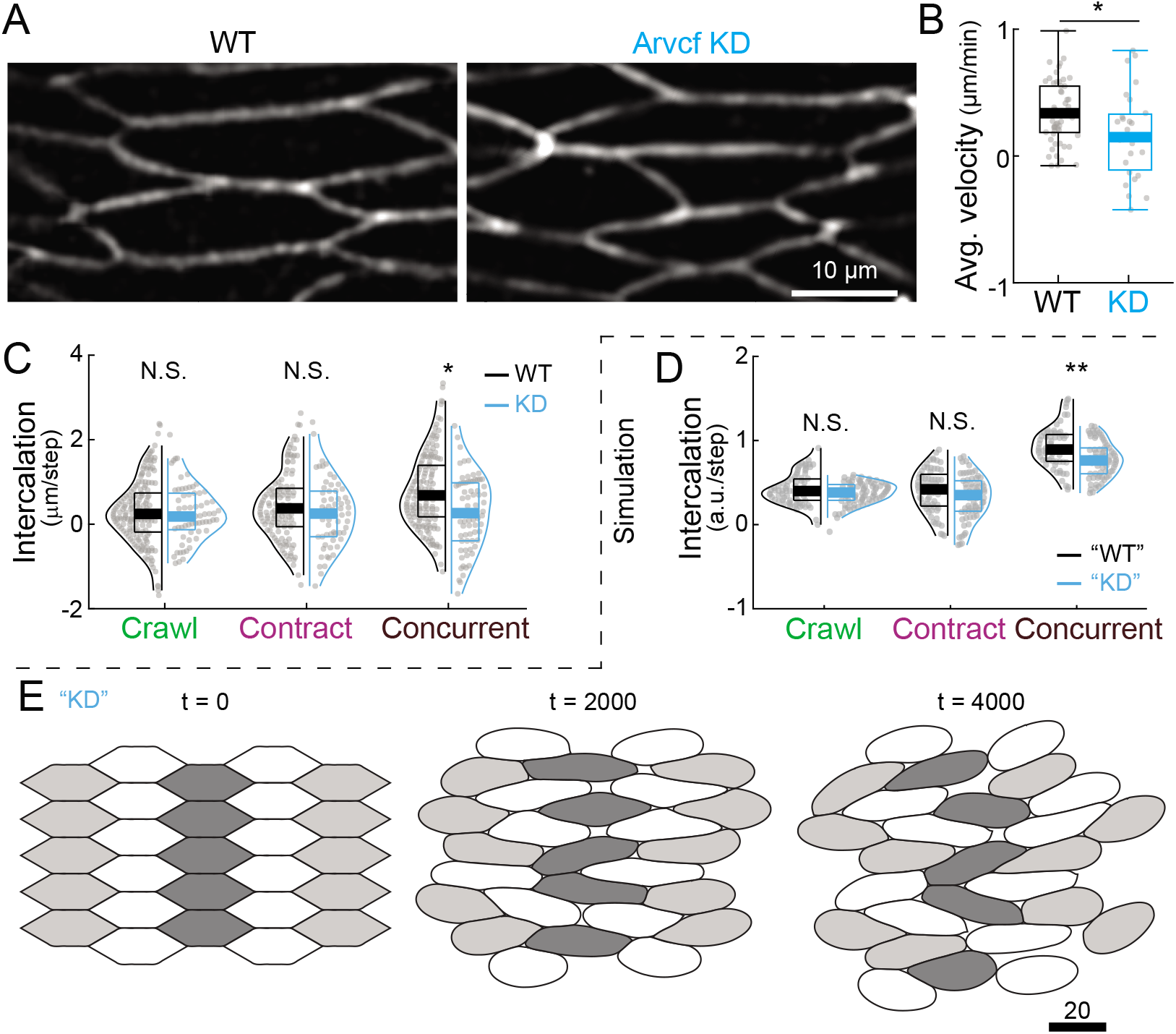
The Arvcf catenin is specifically required for integration of crawling and contraction. (A) Still images of membrane-GFP labeled cells showing both wildtype and Arvcf depleted cells were mediolaterally elongated. (B) Arvcf KD decreased average intercalation velocity. (C) Violin plots of intercalation (vertex displacement) in crawling-only, contraction-only, and concurrent steps. (D) Simulation results recapitulating the concurrent step-specific defect in intercalation. “KD” marks the results using a set of parameters for simulating Arvcf depletion. (E) Representative simulation result of a 27-cell model with parameters simulating Arvcf depletion. Tissue-wide convergent extension is reduced.

Turning to our model, we first considered that loss of Arvcf elicits a small but significant reduction in cortical cadherin levels, resulting in enlarged gaps between cells (Huebner et al., 2021b). Curiously, while reducing adhesion alone in our model could elicit the formation of extracellular gaps (**Fig. 6A, B**), it could not recapitulate the specific disruption of concurrent steps that we observed for Arvcf loss *in vivo* (**Fig. 6C-E**).

This result suggested that Arvcf, plays a more complex role, so we performed a broader survey of parameter space in our model. Strikingly, only one set of parameters in the ranges tested here recapitulated the concurrent step-specific defect in intercalation (**Fig. 7D**). These parameters (−25% adhesion 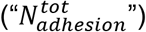; −23% crawling (“*F_crawl_*”); +7% contraction (“*M_pulse_*”)) also recapitulated the defect in amplification of crawling-associated protrusions during concurrent steps (**Supp. Fig. 8E**), leaving the amplification of contraction-associated forces unaltered (**Supp. Fig. 8F**). Finally, these model parameters also recapitulated the failure of convergent extension (**Fig. 7E**). Thus, our new model faithfully modelled *in silico* both fine and gross aspects of the Arvcf knockdown phenotype that we observed *in vivo*.

## Conclusions

Here, we have combined live imaging and a new approach to modeling to explore the mechanisms by which two modes of cell motility – protrusive crawling and cortical contraction – collaborate to drive a crucial morphogenetic process in early vertebrate embryos. Our live imaging approach allowed us to independently identify and quantify the actin assembly associated with crawling or contraction (**Figs. 1–3)**. By doing so, we show that when crawling and contraction occur concurrently during mesenchymal CE in *Xenopus,* actin assembly associated with both mechanisms is amplified (**Fig. 4**). Moreover, this amplification of actin assembly in turn is associated with significantly improved efficacy of cell intercalation. Similar mechanosensitive responses of actomyosin are observed in diverse epithelial cells (e.g. (Fernandez-Gonzalez et al., 2009; Martin et al., 2010), so our work here in mesenchymal cells in *Xenopus* suggests that the crawling- and contraction-based intercalation mechanisms in epithelial cells (e.g. (Sun et al., 2017; Williams et al., 2014) may be similarly coupled.

In addition, our new approach to modeling of CE provides a substantial advance over previous models. It independently captures junction contraction, cell crawling, and cell-cell adhesion, while also treating individual cell cortices independently from one another. The model therefore allows detailed description of both gross tissue movement and also the diverse underlying subcellular behaviors (**Figs. 5, 6)**. This is crucial, since individual cell-cell junctions during *Xenopus* CE display very local heterogeneities in their mechanical properties (Huebner et al., 2021a). Similar local heterogeneity has also been observed during CE in *Drosophila* epithelial cells (Vanderleest et al., 2018) and when junction shortening is artificially induced in cell culture (Cavanaugh et al., 2021). Despite our substantially more complex model, it nonetheless recapitulates both gross and fine characteristics of vertebrate CE observed *in vivo* (**Fig. 5**). Given the model’s ability to provide insights into complex phenotypes observed *in vivo* (**Fig. 7**), we feel this model will provide a very useful resource for the community.

Finally, this work also provides new insights into the role of cell adhesion during CE. The little-studied Arvcf catenin not only tunes cadherin adhesion, but also functionally interacts with the small GTPases RhoA and Rac, negatively regulating RhoA function and conversely promoting Rac function (Fang et al., 2004). How these functional interactions relate to cell behaviors is not known, but our data and modelling suggest that Arvcf-mediated Rac activity is necessary for normal ML crawling, while Arvcf-tempering of RhoA activity is needed to restrain AP contraction. In light of previous data on the cell biological roles of RhoA and Rac during CE (Tahinci and Symes, 2003), a re-examination using mosaic labelling should be highly informative. Thus, our work here provides not only new insights but also new tools for a deeper understanding of *Xenopus* convergent extension specifically and the mechanisms integrating biomechanical forces that drive animal morphogenesis generally.

## Acknowledgements

We thank Dr. Daniel J Dickinson and Dr. Abdul Naseer Malmi-Kakkada for critical reading. This work was supported by grants from the NICHD (R01HD099191;1R21HD103882).

## Supplementary Figure Legends

**Supp. Figure 1.**
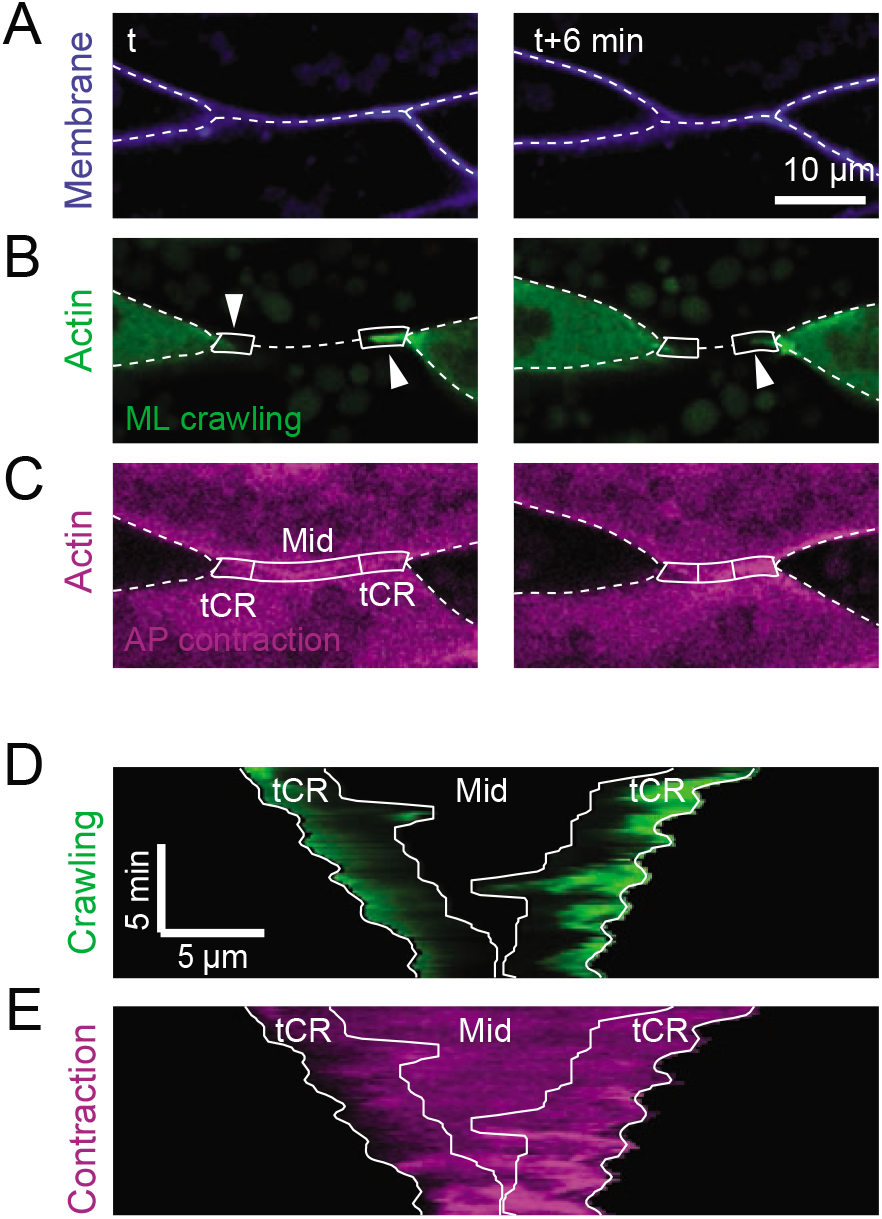
Representative images from a second example. (A-C) Still images from another representative time-lapse movie (**Movie S2**) showing in (A) membrane (blue), in (B) actin in the ML protrusions for crawling (green, arrowheads) and in (C) actin at the AP interface for contraction (purple). Dashed lines mark cell-cell interfaces. Boxes mark tCRs and the middle region (“Mid”). (D, E) Kymograph along the AP cell interface shows spatiotemporal dynamics of actin from the ML protrusions representing the “crawling” signal (D) and actin from the AP cells representing the “contraction” signal (E). White lines outline the tCRs.

**Supp. Figure 2.**
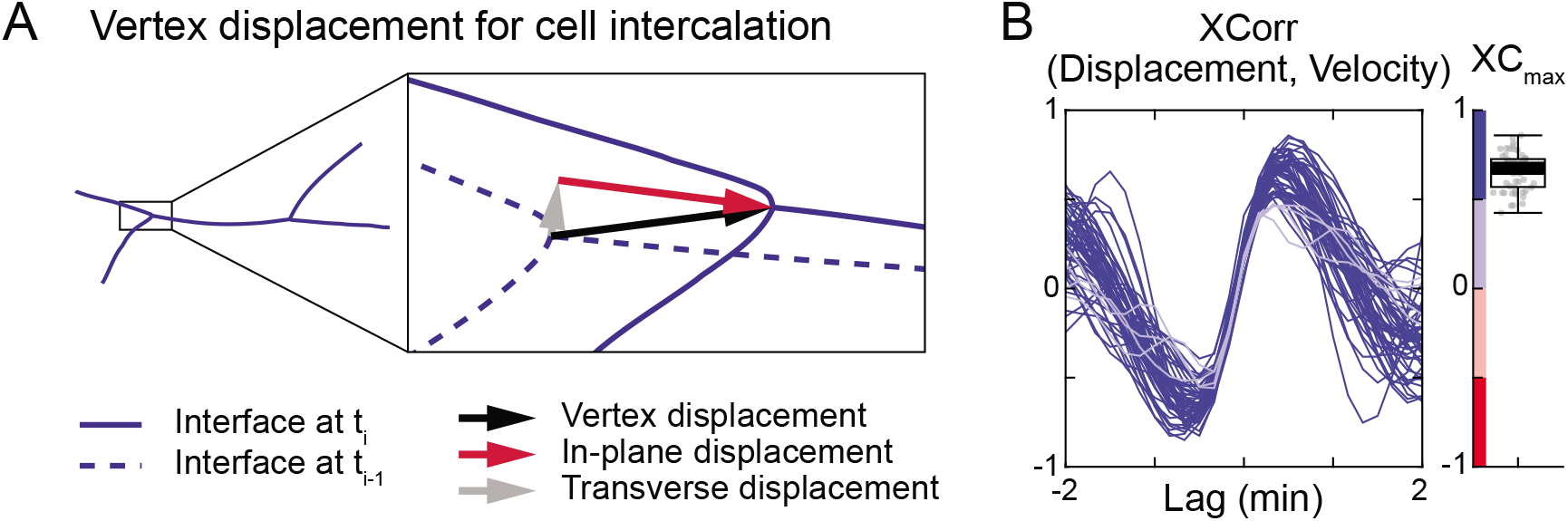
Quantification of cell intercalation. (A) Schematic illustrating the quantification of cell intercalation. Dashed blue lines and solid blue lines are the cell-cell interfaces at two adjacent time points *t_i-1_* and *t_i_*. Black arrow shows the displacement of the tricellular vertex. Gray arrow is the transverse displacement that does not contribute to cell intercalation. Red arrow is the in-plane displacement (i.e., displacement along the AP interface) at t_i_, the only component that contributes to cell intercalation. To keep it simple, the in-plane displacement is referred to as “vertex displacement”, “intercalation displacement”, or “displacement”. (B) Strong cross correlation between intercalation displacement and intercalation velocity, and velocity dynamics consistently leads the dynamics over displacement.

**Supp. Figure. 3.**
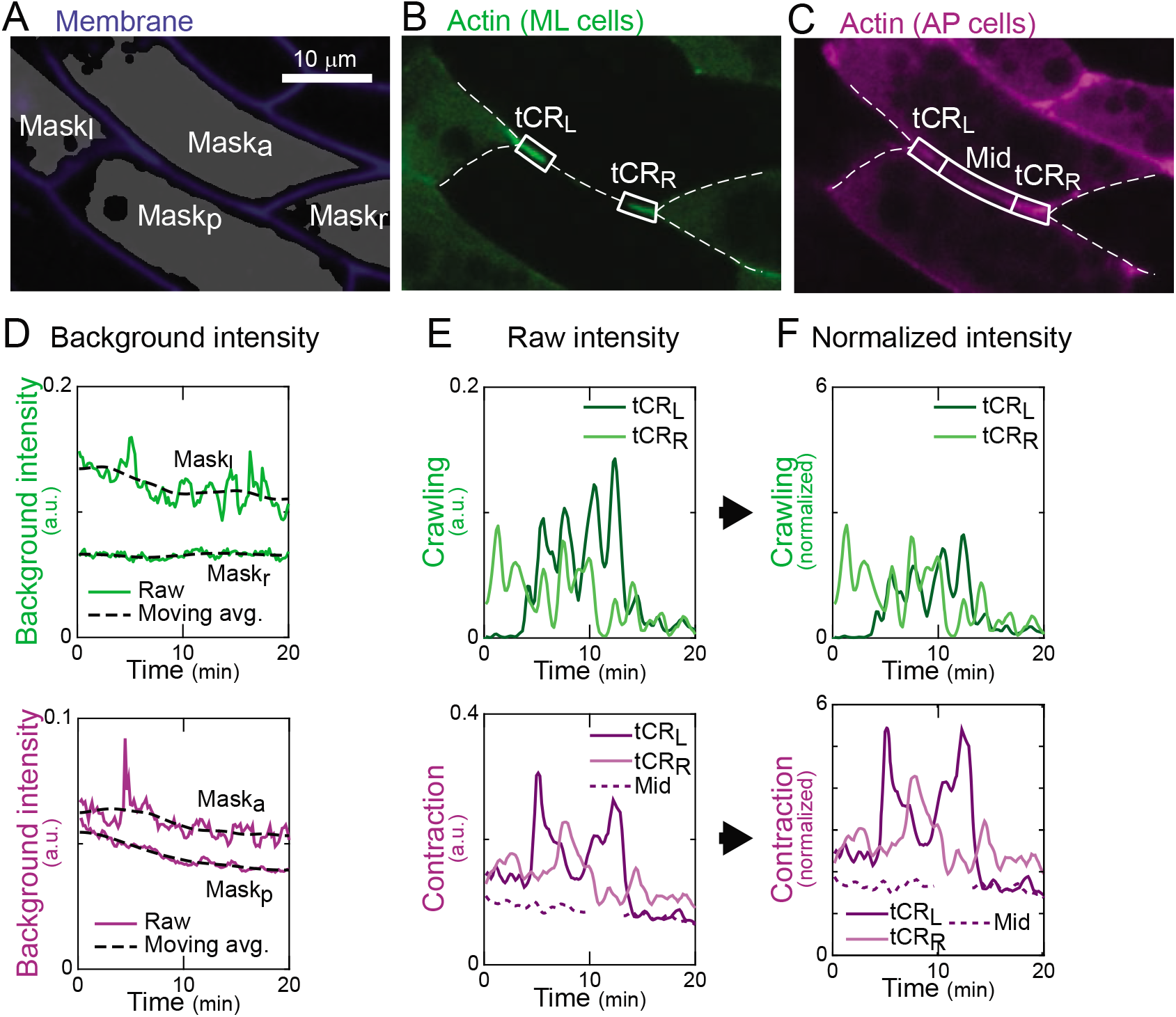
Quantification of actin dynamics for crawling and contraction. (A) Membrane labeling (blue) was used for segmentation and generating cytosolic masks (gray) for quantifying background intensity. (B, C) Actin in the ML cells (B) and in the AP cells (C). Cellcell interfaces were detected using the membrane labeling (white dashed lines). White boxes mark the left tCR (tCR_L_), right tCR (tCR_R_), as well as the mid region (Mid). See Methods for the detailed definition of tCR and Mid regions. (D) Background fluorescence intensity of actin (solid lines) was defined as the mean actin intensity in the cytosolic masks shown in (A). Their moving average over 6 min (black dashed lines) was quantified and used for normalization later. (E) Raw fluorescence intensity of actin for crawling and contraction was defined as the mean actin intensity in the boxed regions in (B) & (C). (F) Fluorescence intensity normalized by the moving average of background intensity in (D).

**Supp. Figure 4.**
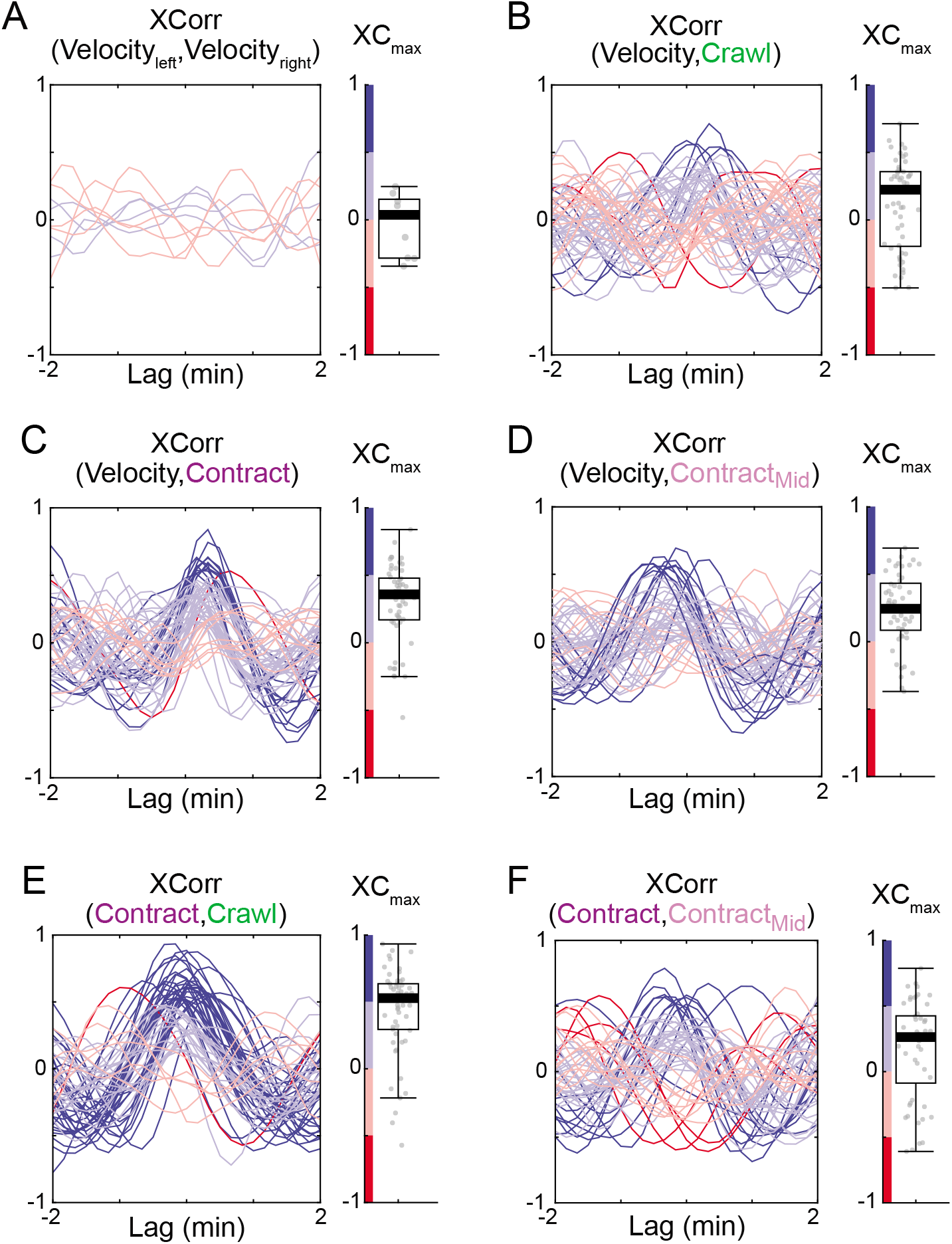
Cross-correlation analysis of cell intercalation velocity and bulk actin dynamics. (A) Low cross-correlation between intercalation velocity from a paired left and right tricellular vertices sharing the same AP interface. n = 9. (B-D) Low cross-correlation between cell intercalation velocity and bulk actin dynamics for crawling (B), tCR contraction (C), and Mid contraction (D). (E) Strong cross-correlation between actin dynamics for crawling and tCR contraction. (F) Low cross-correlation between actin dynamics for tCR contraction and Mid contraction, demonstrating spatial heterogeneity along the AP interface. All show a wide range of correlation with variable lag time. Each line represents cell intercalation and actin dynamics at one tCR. n > 53.

**Supp. Figure 5.**
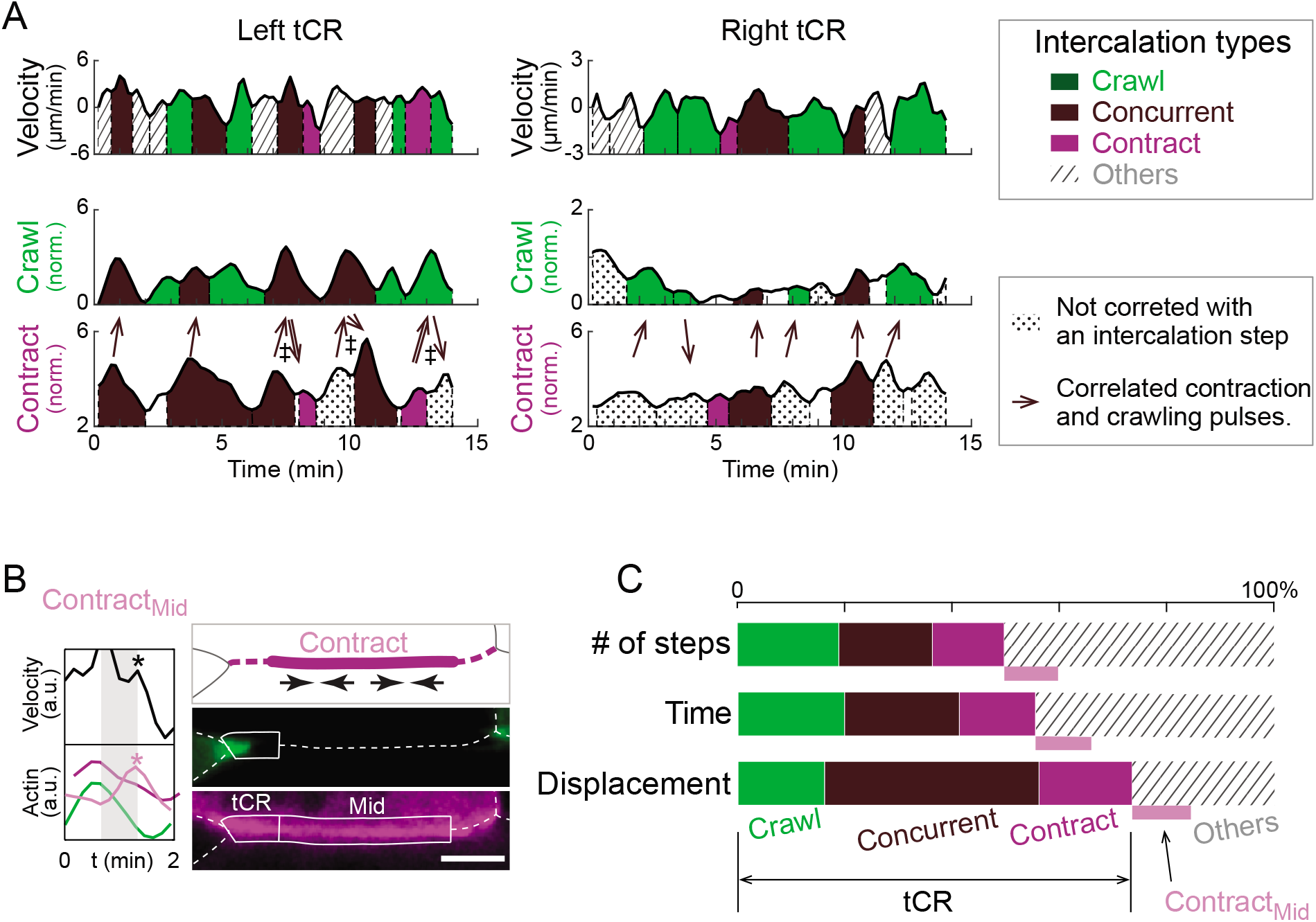
Classification of intercalation steps. (A) Representative intercalation step analysis for the tCRs in **Fig. 1I-M (Movie S1)**. For each tCR, the intercalation velocity, crawling signal, and contraction signal were plotted verses time. Peaks and valleys were detected using customized scripts and individual peaks were separated with dashed lines at the adjacent valleys. An intercalation step was defined as an individual peak on the velocity trace and was classified based on its cross-correlation with pulses on the crawling and/or contraction signals. The correlated peaks were color-coded as indicated. Intercalation steps that had no crosscorrelation with crawling nor contraction pulses were filled with diagonal strips, while contraction and crawling pulses that had no cross-correlation with any intercalation step were filled with dots. Brown arrows label correlated contraction and crawling pulses. Brown double-line arrows mark correlated crawling and contraction pulses that are associated with different velocity peaks. Daggers mark crawling and contraction pulses that were correlated successively. Lagtime window for all cross-correlation analysis was 40 sec, and the threshold for cross-correlation coefficient was 0.5. (B) Examples showing an intercalation step correlated with a contraction pulse from the middle region of an AP interface, and not correlated with actin dynamics in the tCR. See **Movie S7**. (C) **Fig. 2F** replotted in which Mid contraction (“Contract_Mid_”) was also considered. The majority of cell intercalation is attributed to actin dynamics at tCRs and therefore the Mid contraction was neglected in this study.

**Supp. Figure 6.**
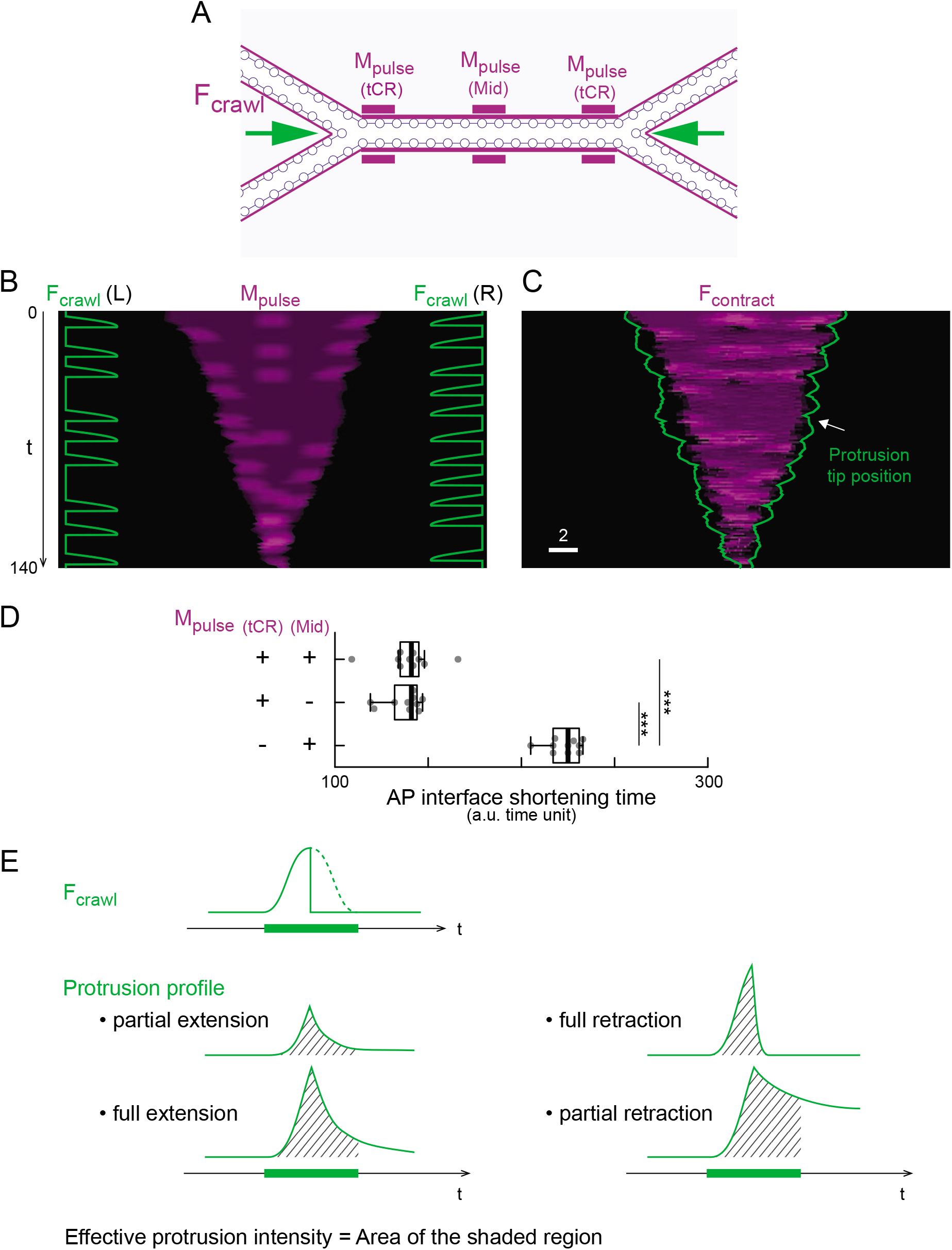
Design of crawling and contraction in a four-cell model. (A) Schematic showing the application of crawling force *F_crawl_* and actomyosin pulses *M_pulse_*. Crawling forces on the left and right cells were applied on the tricellular vertices at randomized timepoints and toward each other exclusively. Actomyosin pulses were applied at both tricellular regions and the middle of the AP interface. (B) Representative simulation input of crawling force and actomyosin pulses. Green traces show the profile of crawling forces on the left and right tricellular vertices. Purple kymograph shows the spatiotemporal dynamics of the applied actomyosin pulses along the shortening AP interface. (C) Simulation results from the input in (B). Purple kymograph shows the calculated contraction forces *F_contract_* along the AP interface. Green traces mark the tip of the protrusions from the left and right cells. (D) Bar plot showing the insignificance of actomyosin pulses from the middle region of an AP interface. (E) Schematic showing the definition of the proxy for protrusion actin assembly. Depending on the cell-cell interaction, protrusion profile varies with the same applied crawling force. The integral of protrusion length over time, defined as the effective protrusion intensity, was used as a proxy for the protrusion actin assembly dynamics.

**Supp. Figure 7.**
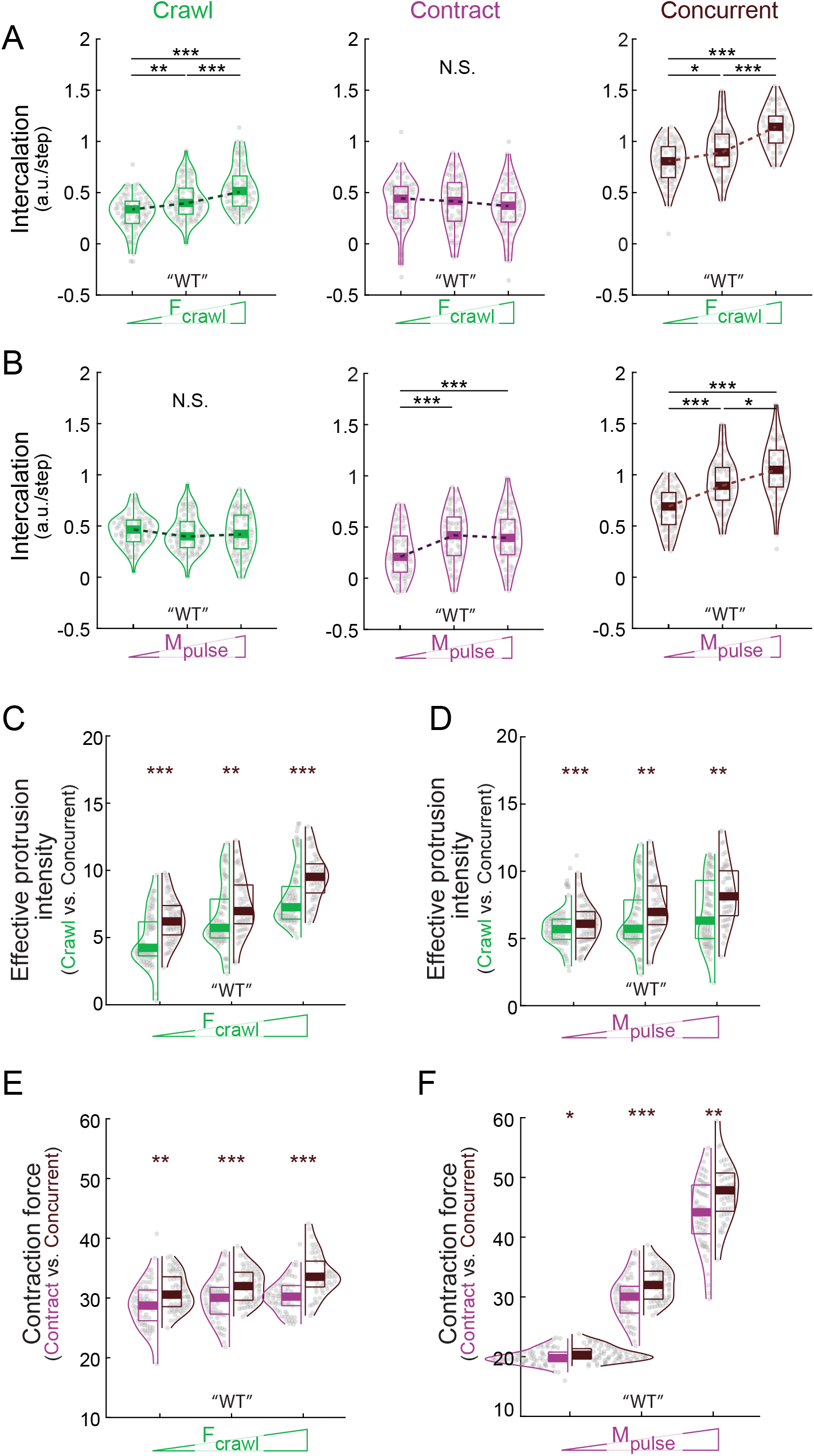
Biomechanical synergy of crawling contraction integration over a wide range of crawling forces and contraction pulses in the *in-silico* model. (A, B) Effect of increasing crawling force (A) or increasing actomyosin pulses (B) on intercalation (vertex displacement) during crawling only, contraction only, and concurrent steps in the model. (C, D) Synergistic effect of concurrent crawling and contraction on the protrusion dynamics over a wide range of crawling forces and actomyosin pulses. (E, F) Synergistic effect of concurrent crawling and contraction on the contraction force over a wide range of crawling forces and actomyosin pulses. “WT” marks the values used for simulating the wildtype condition.

**Supp. Figure 8.**
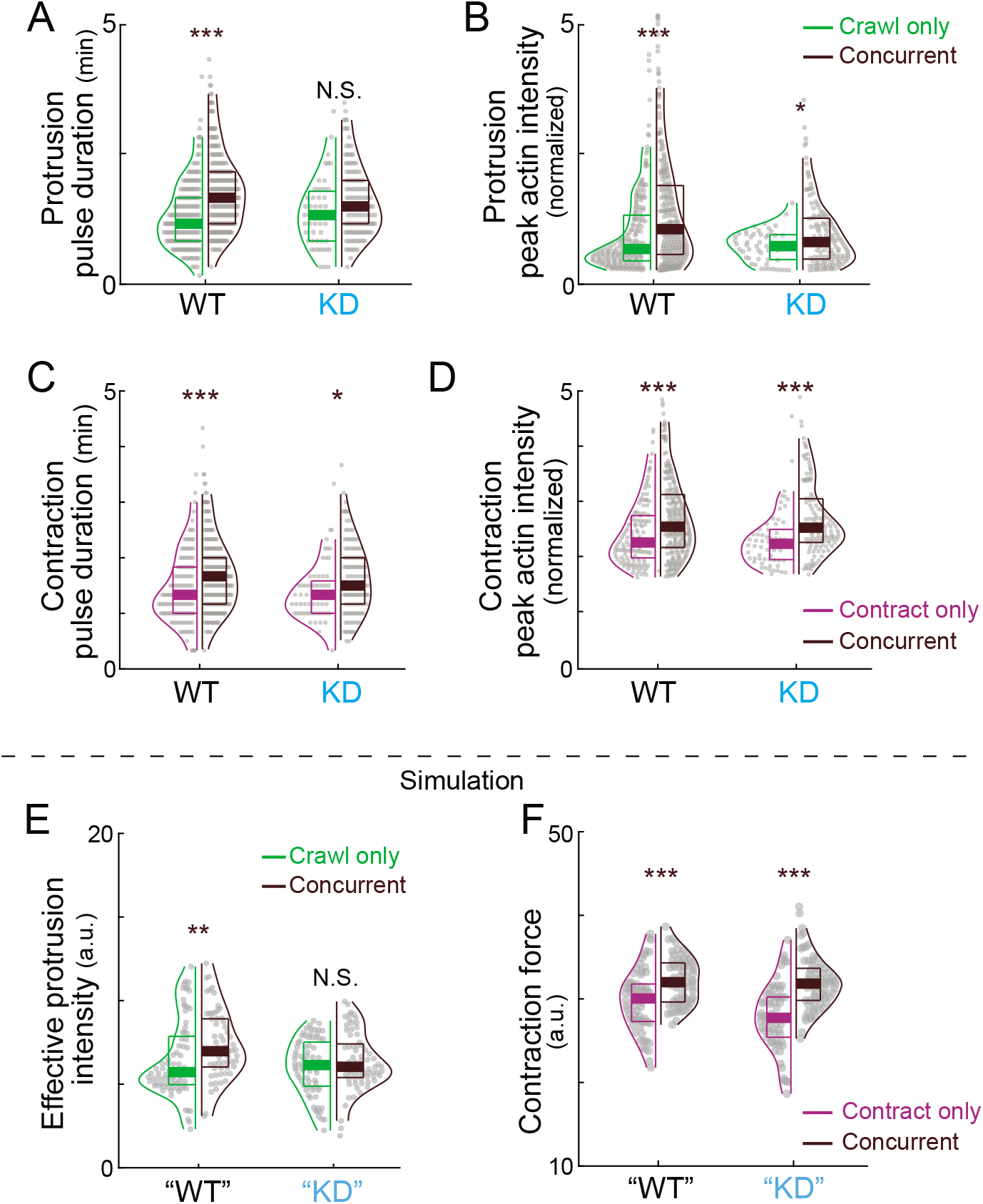
Supplementary results of Arvcf knockdown. (A) Violin plots showing that elongated protrusion duration during concurrent steps normally observed was disrupted in Arvcf depleted cells. (B) Violin plots showing that amplified protrusion assembly during concurrent steps normally observed was diminished in Arvcf depleted cells. (C) Violin plots showing that elongated contraction pulse duration during concurrent steps was not altered in Arvcf depleted cells. (D) Violin plots showing that amplified contraction pulses during concurrent steps was not altered in Arvcf depleted cells. (E) Modeling results showing diminished amplification of the effective protrusion intensity during concurrent steps with the “KD” parameters. (F) Modeling results showing no difference on the amplification of contraction force during concurrent steps with the “KD” parameters.

## Supplementary table legend

**Supp. Table 1.** Parameters for the CE vertex model. Asterisks mark values used to simulate the wildtype condition, daggers for the low adhesion condition, and section signs for the Arvcf knockdown.

| Parameter |  |  | Value | Reference |
| --- | --- | --- | --- | --- |
| Iteration | Viscosity | $\eta$ | 20 | - |
| | Timestep | $\Delta t$ | 0.05 | - |
| Contraction | Stall force | $f_{stall}$ | 4 | (Mitrossilis et al., 2009b) |
| | Actomyosin density | $M_{base}$ | 2 (AP)<br>1 (ML) | (Mitrossilis et al., 2009a) |
| | | $M_{pulse}$ | 2, 4.5*, 4.8<sup>§</sup>,<br>7 | |
| | Min. strain rate | $\varepsilon_o$ | 0.8 | (Mitrossilis et al., 2009a) |
| Protrusion | Lamellipodia force | $F_l$ | 1 | - |
| | Lamellipodia duration | $T_l$ | 100 | - |
| | Protrusion force | $F_p$ | 14<sup>§</sup>, 18*, 22 | (Abraham et al., 1999; Prass et al., 2006) |
| | Protrusion duration | $T_p$ | 6 | - |
| Adhesion | Force scale | $f^*$ | 0.005 | (Buckley et al., 2014) |
| | Catch unbinding rate | $\phi_c$ | 1.0322 | (Buckley et al., 2014) |
| | Slip unbinding rate | $\phi_s$ | 2.9671 | (Buckley et al., 2014) |
| | Rebinding rate | $\gamma$ | 2 | - |
| | Total available adhesion unit | $N_{adhesion}^{tot}$ | 600<sup>†</sup>, 900<sup>§</sup>,<br>1200*, 1500 | (Truong Quang et al., 2013) |
|  |  | AP/ML | 1:0.6 | unpublished data |
| | Adhesion range | $d_a$ | 0.25 | - |
| Cytoplasm | Rest area/Initial area | $A_r/A_0$ | 1.002 | (Petrie et al., 2014) |
| | Stiffness | $k_C$ | 200 | (Petrie et al., 2014) |
| Repulsion | Maximum force | $F_{rmax}$ | 6 | - |
| | Force at zero distance | $F_{r0}$ | 2 | - |
| | Repulsion range | $d_r$ | 0.025 | - |

## Supplementary Movie legend

**Movie S1. Representative timelapse movie showing the mosaic labeling to distinguish crawling and contraction signals for cell intercalation.** Cells were uniformly labeled with the membrane marker memBFP and mosaically labeled for different colors of an actin biosensor (Lifeact-RFP/Lifeact-GFP). The uniform membrane-BFP (blue) was used to segment cells (white dashed lines), and Lifeact-GFP and -RFP used to unambiguously quantify different populations of actin (green for ML cells and purple for AP cells). Still frames and kymograph along the AP cell interface were shown in **Fig. 1 E-H**.

**Movie S2. Another representative timelapse movie showing the mosaic labeling to distinguish crawling and contraction signals for cell intercalation.** Cells were uniformly labeled with the membrane marker memBFP and mosaically labeled for different colors of an actin biosensor (Lifeact-RFP/Lifeact-GFP). The uniform membrane-BFP (blue) was used to segment cells (white dashed lines), and Lifeact-GFP and -RFP used to unambiguously quantify different populations of actin (green for ML cells and purple for AP cells). Still frames and kymograph along the AP cell interface were shown in **Supp. Fig. 1**.

**Movie S3. Pulsatile nature of cell intercalation movement and actin dynamic for crawling and contraction.** Left panel: Same as Movie S1 with dashed box tracing the left tricellular region (tCR) for analysis. White arrows are shown when the crawling protrusion (top) or contraction signal (bottom) peaks. Right panel: Traces of intercalation velocity (i.e. left vertex displacement; top) and traces of crawling (green) and contraction (purple) signals (bottom). Gray dashed line moves over time. Green arrows and purple arrows are shown when crawling signal and contraction signal peak, respectively.

**Movie S4. Crawling step of cell intercalation.** Left panel: traces of intercalation velocity (top) and actin dynamics (bottom) for crawling (green) and contraction (purple). Asterisk marks the timepoint when intercalation velocity peaks. Black arrow marks the velocity peak. Gray boxes mark the time window for correlational analysis. Green arrow marks the crawling peak correlated with the velocity peak. Right panel: Timelapse movie trimmed from **Movie S1**. Dashed box marks the tricellular region. Green arrow traces the crawling peak correlated with the velocity peak.

**Movie S5. Contraction step of cell intercalation.** Left panel: traces of intercalation velocity (top) and actin dynamics (bottom) for crawling (green) and contraction (purple). Asterisk marks the timepoint when intercalation velocity peaks. Black arrow marks the velocity peak. Gray boxes mark the time window for correlational analysis. Purple arrow marks the contraction peak correlated with the velocity peak. Right panel: Timelapse movie trimmed from **Movie S1**. Dashed box marks the tricellular region. Purple arrow traces the contraction peak correlated with the velocity peak.

**Movie S6. Concurrent crawling and contraction step of cell intercalation in which crawling leads.** Left panel: traces of intercalation velocity (top) and actin dynamics (bottom) for crawling (green) and contraction (purple). Asterisk marks the timepoint when intercalation velocity peaks. Black arrow marks the velocity peak. Gray boxes mark the time window for correlational analysis. Green arrow marks the crawling peak correlated with the velocity peak, while the purple arrow for the contraction peak. Right panel: Timelapse movie trimmed from **Movie S1**. Dashed box marks the tricellular region. Green arrow traces the crawling peak correlated with the velocity peak, and purple arrow for the contraction peak.

**Movie S7. Concurrent crawling and contraction step of cell intercalation in which contraction leads.** Left panel: traces of intercalation velocity (top) and actin dynamics (bottom) for crawling (green) and contraction (purple). Asterisk marks the timepoint when intercalation velocity peaks. Black arrow marks the velocity peak. Gray boxes mark the time window for correlational analysis. Green arrow marks the crawling peak correlated with the velocity peak, while the purple arrow for the contraction peak. Right panel: Timelapse movie trimmed from **Movie S1**. Dashed box marks the tricellular region. Green arrow traces the crawling peak correlated with the velocity peak, and purple arrow for the contraction peak.

**Movie S8. Contraction step in which contraction comes from the middle of an AP interface.** Left panel: traces of intercalation velocity (top) and actin dynamics (bottom) for crawling (green) and contraction (purple from tCR and pink from the middle region of the AP interface). Asterisk marks the timepoint when intercalation velocity peaks. Black arrow marks the velocity peak. Gray boxes mark the time window for correlational analysis. Pink arrow marks the contraction peak correlated with the velocity peak. Right panel: Timelapse movie trimmed from **Movie S1**. Dashed box marks the tricellular region. Pink arrow traces the contraction peak from the middle of the AP interface which is correlated with the velocity peak.

**Movie S9. Four-cell model of convergent extension.** Representative result from a four-cell model simulating cell intercalation and convergent extension between four cells. Medially-directed crawling forces (“*F_crawl_*”) and heterogenous actomyosin pulses along the AP cell interface (“*M_pulse_*”) were applied at randomized timepoints. AP cell interface shortened completed followed by the formation and elongation of a new AP directed cell-cell interface.

**Movie S10. 27-cell model of convergent extension.** Representative results from a 27-cell model simulating tissue-wide convergent extension. Actomyosin pulses as well as both medially and laterally directed crawling forces were applied stochastically. Multiple rounds of cell intercalation were observed, and the cell cluster converged and extended.

## Methods

### *Xenopus* embryo manipulations

Ovulation was induced by injecting adult female Xenopus laevis with 600 units of human chorionic gonadotropin (HCG, MERCK Animal Health) and animals were kept at 16 dc overnight. Eggs were acquired the following day by squeezing the ovulating females and eggs were fertilized *in vitro.* Eggs were dejellied in 2.5% cysteine (pH 7.9) 1.5 hours after fertilization and reared in 1/3x Marc’s modified Ringer’s (MMR) solution. For micro-injection, embryos were placed in 2% ficoll in 1/3x MMR during injection and washed in 1/3x MMR 30 min after injection. Embryos were injected in the dorsal blastomeres at the 4-cell stage targeting the C1 cell at 32-cell stage and presumptive notochord. Keller explants were dissected at stage 10.25 in Steinberg’s solution using hair tools.

### Plasmids and Morpholinos

The Arvcf morpholino has been previously described (5’-ACACTGGCAGACCTGAGCCTATGGC-3’ (Fang et al., 2004) and was ordered from Gene Tools, LLC. Lifeact-GFP and Lifeact-RFP were made in pCS and membrane-BFP in pCS2.

### mRNA and morpholino microinjections

Capped mRNAs were generated using the ThermoFisher SP6 mMessage mMachine kit (Catalog number: AM1340). mRNAs were injected at the following concentrations per blastomere: Mem-BFP (75 pg), Lifeact-RFP (75 pg), and Lifeact-GFP (75 pg). Arvcf morpholino was injected at a concentration of 30ng per blastomere.

### Imaging *Xenopus* explants

Explants were submerged in Steinberg’s solutions and cultured on glass coverslips coated with Fibronectin (Sigma-Aldrich, F1141) at 5 μg/cm^2^. After 5-hour incubation at room temperature, we started standard confocal time-lapse imaging using a Nikon A1R. Images of membrane-BFP, Lifeact-GFP, and Lifeact-RFP were taken at a focal plane 5 μm deep into the explant and at an interval of 10 sec.

### Cell segmentation

All image process and analysis were performed using customized MATLAB scripts if not mentioned otherwise. We first performed cell segmentation and junction detection on images of Membrane-BFP. Briefly, we used pixel classification in ilastik (Berg et al., 2019), a machine-learning based open-resource image analysis tool, to classify pixels at cell-cell interface and pixels in cytoplasm. This process converted a time-lapse movie of fluorescence intensity images to a time-lapse movie of probability images for cell-cell interface. We then extracted skeletons where the probability peaks at each time point for a robust detection of cellcell interfaces and cell segmentation.

### Kymograph preparation

We customized the preparation of kymograph so that it not only displays actin fluorescence intensity values from crawling (ML cells) or contraction (AP cells) signals but also displays the movement of tricellular vertices explicitly. Briefly, we extracted 2 μm thick bands along the AP cell interface from images of actin labeling, linearized the bands, calculated the normalized mean fluorescence intensity across the thickness, then stacked data from each timepoint along the time axis. For a direct display of tricellular vertex movement, we used in-plane displacement at each time point, defined in **Supp. Fig. 2A**, as a reference to align the kymograph.

### Measurement of cell intercalation

We defined cell intercalation as the displacement of a tricellular vertex along the direction to shorten the AP interface (**Supp. Fig. 2A**). Each tricellular vertex was tracked over time on the segmented images and its displacement in the tangential direction of the AP interface (“in-plane”) between adjacent timepoints was quantified (**Supp. Fig.2A**). The in-plane displacement is the only component that contributes to cell intercalation and junction shortening. The transverse movement of a tricellular vertex causes junction rotation and was neglected here. To keep it simple, the in-plane displacement is referred to as “vertex displacement”, “intercalation displacement”, or “displacement”. We quantified cell intercalation at each tCR separately because their behaviors were independent (**Supp. Fig. 4A**).

### Measurement of actin dynamics for crawling and contraction

For the quantification of actin intensity associated with crawling or contraction, a 2 μm thick band along the AP cell interface was extracted from the fluorescent image of actin for ML cells or AP cells respectively (**Supp. Fig. 3B, C**). For each time point, we divided such bands into tricellular regions (tCRs) and the middle region (Mid) for the analysis of regional actin dynamics. The tCRs have a minimum length of 4 μm at any time point and covered the entire protrusions from the ML cells if there were any. The middle regions were complementary to the tCRs and had a minimum length of 2 μm (**Supp. Fig. 3C**). The raw crawling and contraction signals for cell intercalation were the mean actin fluorescence intensity in the correspondence tCRs or the middle region as specified (**Supp. Fig. 3E**). The necessity for such a regional analysis is apparent, since actin dynamics are highly heterogeneous long the AP interface (**Fig. 1L, M**). The low correlation between the contraction signal in a tCR and that in the adjacent middle region also supports this regional analysis (**Supp. Fig. 4F**). Since contraction from the middle region of a junction has minor contribution to cell intercalation (**Supp. Fig. 5C**), this manuscript focuses on actin dynamics in tCRs and contraction signals refer to contraction in tCRs if not otherwise specified.

### Normalization of fluorescence intensity signals

To allow for comparison of fluorescence intensity from different cells and from different actin biosensors, we normalized the fluorescence intensity of each cell with its mean cytosolic background intensity. Briefly, we used cell segmentation mentioned earlier to generate a cytosolic mask for each cell which excluded cellcell interface and cell cortex with a 3 μm margin (**Supp. Fig. 3A**). The mean fluorescence intensity within each cytosolic mask was quantified and its moving average over 6 min was calculated and used for normalization (**Supp. Fig. 4D-F**).

### Intercalation step analysis by correlating actin dynamics and cell movement

To understand the underlying mechanisms for cell intercalation, we first performed cross-correlational analysis between the intercalation velocity traces and the traces of actin dynamics for crawling or contraction (**Supp. Fig. 4B-D**). Vertex velocity traces were chosen rather than the vertex displacement for the dynamics of cell intercalation, because displacement is the integral of a velocity and trails the dynamics of the velocity (**Supp. Fig. 3B**). The bulk analysis shown low cross correlation between the cell movement and actin dynamics (**Supp. Fig. 4B-D**), suggesting a more complex mode of cell intercalation than crawling or contraction alone. Meanwhile, the contraction signals and crawling signals in tCRs were better correlated in over half of the cases (**Supp. Fig. 4E**), suggesting an active integration between them.

To infer the driving force for cell intercalation at a finer temporal resolution, we took the advantage of the pulsatile nature of both actin dynamics and cell movement during CE. We first identified peaks in the intercalation velocity curves as single intercalation “steps”, then searched for cross-correlated peaks in the crawling and contraction signals from tCRs within a 40-sec window preceding the velocity peak (**Fig. 2A**; e.g. **Supp. Fig. 5A**; threshold for crosscorrelation: 0.5). If an intercalation step is correlated with a peak in the crawling or contraction signal, it is categorized as a crawling or contraction intercalation step (**Fig. 2B**, **Movie S4**; **Fig. 2C**, **Movie S5**). If an intercalation step is correlated with a pair of contraction and crawling peaks, it is categorized as a concurrent intercalation step (**Fig. 2D, E; Movie S6, 7**). Within the 40-sec window, we observed comparable number of cases when a crawling peak precedes a contraction peak (**Fig. 2D**) and cases when a contraction peak precedes a crawling peak (**Fig. 2E**). Besides the crawling, contraction, and concurrent intercalation steps, which are the major players, we also observed intercalation steps that are correlated with contraction peaks from the middle region of an AP interface (**Supp. Fig. 5B**; **Movie S8**). Compare with actin dynamics in tCRs, contraction in the middle region has much less contribution to cell intercalation (**Supp. Fig. 5C**).

### Statistical analysis

For intercalation analysis, 57 tricellular vertices from wildtype cells from at least 10 independent experiments and 27 tricellular vertices from Arvcf KD cells from at least 3 independent experiments are pooled. For step analysis results from four-cell models simulating shortening of AP interface (as in **Fig. 5I-L**; **Fig. 6C-G**; **Fig. 7D**; **Supp. Fig. 7**; and **Supp. Fig. 8E, F**), 19 independent simulations were run for each set of parameters. In box plots, each dot represents a datapoint from a vertex. The central line is the median, the box extends vertically between the 25^th^ and 75^th^ the percentiles, and the whiskers extend to the lower and upper limits that do not include the outliers. For violin plots and split violin plots, each dot represents an individual intercalation step/event (as in **Fig. 3**, **Fig. 5I, J**, **Fig. 6C-E** and **Fig. 7C, D**), or an individual crawling or contraction pulse (as in **Fig. 4**, **Fig. 5K, L**, **Fig. 6F, G**, and **Supp. Fig. 7**) identified from the data pool. Besides the included boxes, the violin plots also show the probability density extending from the lower limits to the upper limits that do not include the outliers. All P values are calculated using Wilcoxon Rank Sum Test (A.K.A Mann Whitney U Test; MATLAB statistics toolbox). *p<0.05; **p<0.005; ***p<0.0005; N.S., not significant.

### Vertex model to simulate convergent extension

To study how crawling and contraction act in concert to drive convergent extension, we designed a novel cell-based CE model that offers a detailed description of subcellular behaviors and allows for cells taking an arbitrary shape under subcellular forces. The design of the model is inspired by a cell-based model for the gastrulation of Nematostella vectensis (Tamulonis et al., 2011). Each cell is initially a hexagon and is represented by a 90-vertex polygon (**Fig. 5A**). Interfaces between any neighboring cells are allowed to connect via cell-cell adhesion (**Fig. 5B, C**).

The dynamics of the model is driven by simple Newtonian mechanics with the basic assumptions: (1) vertices are embedded in viscous medium that applies a viscous dragging force with the damping parameter *η* and (2) inertia vanishes. This leads to the governing equation for the evolution of the position ***x**_i_* of vertex *i* determined by

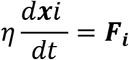

where ***F_i_*** is the total force acting on the vertex *i*. Each simulation step is run by determining the forces acting on each vertex and solving the system of first-order differential equations using two-step Adam-Bashforth method with timestep Δ*t* = 0.05. For each cell, vertices are reset every unit time so that each segment has relatively the same length and regions with high contraction do not have vertices over-crowded.

Inspired by the experimental observations, we considered three key force components in our model: (i) pushing force on the “leading” edge simulating lamellipodia and filopodia-like protrusions (“*F_crawl_*”, **Fig. 5D**), (ii) contractile force on cell edges simulating contraction at cell cortex (“*F_contract_*”, **Fig. 5E**), and (iii) adhesive force between cells simulating cadherin dependent cell adhesion (“*F_adhesion_*”, **Fig. 5C**). Secondary force components incorporated into our model include (iv) elastic cytosolic pressure maintaining the area of a cell and (v) repulsive force between cells to avoid cell collision. Without loss of generosity, this model is a dimensionless, but parameters for the key force components are estimated from experimental data, maintaining their relative values in a physiological relevant range (see **Supp. Table.1**).

#### (i) Pushing force for cell crawling and protrusion extension

We considered two force components associated with the crawling force, *F_crawl_*, one for lamellipodia-like pushing on the “leading” edge and one for filopodia-like pushing at a more focused region. The “leading” edge forms stochastically and polarity-dependently (i.e., randomly toward median or lateral for convergent extension). Small pushing forces with a cosinusoidal profile and the maximum magnitude of *F_l_* are applied on vertices along the leading edge for a period of *T_l_*. The cosinusoidal force profile is initially centered around the anteroposterior center of the cell and could be shifted toward a protrusion when one forms as described below. Filopodia-like protrusions form stochastically on vertices in the tricellular regions on the “leading” edges. When a protrusion site is identified, a quartile-sine force pulse with the magnitude of *F_p_* and the duration of *T_p_* is applied to the vertex in the direction toward the interface between its two neighboring cells (**Supp. Fig. 6A**). For the analysis of different intercalation types, we simulated the shortening of AP interfaces exclusively and set the filopodial protrusions forming at defined time points (**Fig. 5H**), so that we can distinguish crawling-only intercalation events from concurrent events.

#### (ii) Contraction at cell cortex

Contraction at cell cortex is modeled as pieces of Hill’s muscle connected in series. The contractile force between two neighboring vertices equals:

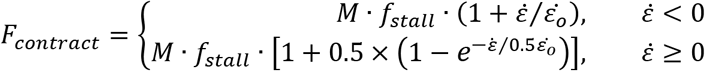

where *M* is the density of actomyosin, *f_stall_* is the stall force, 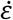 is the strain rate in each segment defined by adjacent vertices, and 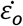 is the maximum contractile strain rate when force vanishes. To mimic actomyosin pulses long the AP interface (**Fig. 1M**), *M* is composed of a polarity-dependent baseline of actomyosin *M_base_*, plus extra half-sine pulses *M_pulse_* in the tricellular regions and/or the middle of the AP interface for a period of *T_m_* (**Supp. Fig. 6A, B**). The extra actomyosin pulses form stochastically in the 4-cell or 27-cell models. However, for the analysis of different intercalation types, we simulated the shortening of AP interfaces exclusively and set the actomyosin pulses at defined time points in tCRs only (**Fig. 5H**), so that we can distinguish contraction-only intercalation events from concurrent events.

#### (iii) Cell-cell adhesion and force transmission

Cell-cell adhesion is based on cadherin clustering via its trans- and cis-interaction, and depends on force transmission via cadherin clusters and cadherin-catenin complex binding to the actomyosin network (Lecuit and Yap, 2015). We simplified cell adhesion as adhesion clusters of the size of *N_adhesion_* binding cell edges of adjacent cells, while holding an adhesive force of *F_adhesion_* (**Fig. 5C**). The dynamics of adhesion clustering and the adhesion force are interdependent and we used a catch-bond model for the simulation (Buckley et al., 2014; Novikova and Storm, 2013; Rakshit et al., 2012).

Briefly, the force dependent unbinding rate for a single catch-bond can be expressed as *k_u_*(*f*) = *k*_0_ · [*exp*(–*f*/*f*^*^ + *φ_c_*) + exp (*f*/*f*^*^ – *φ_s_*)], where *k*_0_ is a reference unbinding rate and set to 1 s^-1^, *f* is the force on the bond, *f*^*^ is a force scale used to non-dimensionalize all forces, and *φ_c_* and *φ_s_* represent zero-force unbinding rates associated with the catch and slip portion of the bond dynamics, respectively. Considering an adhesion cluster of the size *N_adhesion_* and a tensile load *F_adhesion_* uniformly distributed on all bonds, the temporal evolution of the cluster size is expressed as

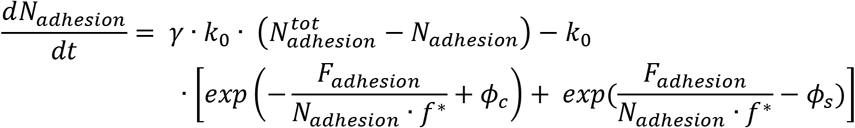

where *γ* is a dimensionless rebinding rate and 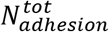 is the polarity-dependent total adhesion units available in the vicinity. Under a quasistatic state, *N_adhesion_* and *F_adhesion_* can reach their maximum given by 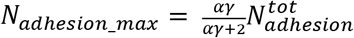 and *F_adhesion_max_* = *N_adhesion_max_* · *Φ_max_* · *f*^*^, where *α* = *exp* (*φ_s_*/2 – *φ_c_*/2) and *φ_max_* = (*φ_s_* + *φ_c_*)/2 (see detailed deduction in (Novikova and Storm, 2013).

Clustering is designed to initiate between any vertex-edge pair from two neighboring cells when the distance between them, *d* is larger than *d_r_*, a repulsion limit defined later to avoid cell collision, but smaller than *d_a_*, the adhesion limit. The two anchoring points of the cluster can move as the cell deforms and can sit anywhere on cell edges.

At the end of each iteration timestep, the adhesive force applied via an adhesion cluster is estimated by *F_adhesion_e_* = *F_a_* · *N_adhesion_* · (*d* – *d_r_*), where *k_a_* is the spring constant for a bond unit. *k_a_* is estimated assuming a cluster under a slowly increasing tension reaches its maximum size of *N_adhesion_max_* and maximum load of *F_adhesion_max_* at *d_a_*, and it equals *k_a_* = *f*^*^*φ_max_*/*d_a_*.

#### (iv) Cytosolic pressure

Cytoplasm is assumed to be linearly elastic, so the difference between the current cell area *A* and the rest area *A_r_* provides a pressure on cell boundary to maintain the cell size. The cytoplasm potential energy is given by 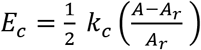, where *k_c_* is the cytoplasm stiffness, and the force on each vertex *i* can be expressed as

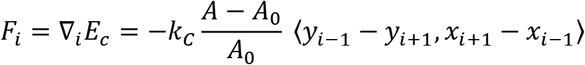

#### (v) Contact repulsion

Repulsive forces are applied to any vertex-edge pair from two neighboring cells if the distance between them *d* if less than *d_r_* or if the vertex is inside the neighboring cell (*d* < 0). The magnitude of this repulsive force is an exponential function of *d* and is given by

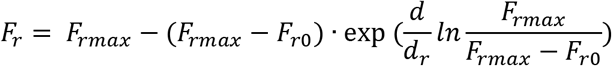

where *F*_*r*0_ is the repulsion at *d* = 0, and *F_rmax_* is the maximum repulsion if the vertex is inside a neighboring cell.

## References

Alt, S., P. Ganguly, and G. Salbreux. 2017. Vertex models: from cell mechanics to tissue morphogenesis. Philos Trans R Soc Lond B Biol Sci. 372.

Becker, S.F.S., R. Mayor, and J. Kashef. 2014. Cadherin-11 Mediates Contact Inhibition of Locomotion during Xenopus Neural Crest Cell Migration. PLOS ONE. 8:e85717.

Belmonte, J.M., M.H. Swat, and J.A. Glazier. 2016. Filopodial-Tension Model of Convergent-Extension of Tissues. PLOS Computational Biology. 12:e1004952.

Berg, S., D. Kutra, T. Kroeger, C.N. Straehle, B.X. Kausler, C. Haubold, M. Schiegg, J. Ales, T. Beier, M. Rudy, K. Eren, J.I. Cervantes, B. Xu, F. Beuttenmueller, A. Wolny, C. Zhang, U. Koethe, F.A. Hamprecht, and A. Kreshuk. 2019. ilastik: interactive machine learning for (bio)image analysis. Nat Methods. 16:1226–1232.

Bertet, C., L. Sulak, and T. Lecuit. 2004. Myosin-dependent junction remodelling controls planar cell intercalation and axis elongation. Nature. 429:667–671.

Brieher, W.M., and B.M. Gumbiner. 1994. Regulation of C-cadherin function during activin induced morphogenesis of Xenopus animal caps. J Cell Biol. 126:519–527.

Brodland, G.W. 2006. Do lamellipodia have the mechanical capacity to drive convergent extension? Int J Dev Biol. 50:151–155.

Buckley, C.D., J. Tan, K.L. Anderson, D. Hanein, N. Volkmann, W.I. Weis, W.J. Nelson, and A.R. Dunn. 2014. The minimal cadherin-catenin complex binds to actin filaments under force. Science. 346:1254211.

Butler, M.T., and J.B. Wallingford. 2017. Planar cell polarity in development and disease. Nature Reviews Molecular Cell Biology. 18:375–388.

Cavanaugh, K.E., M. Staddon, T.A. Chmiel, R. Harmon, S. Budnar, A.S. Yap, S. Banerjee, and M.L. Gardel. 2021. Asymmetric Contraction of Adherens Junctions arises through RhoA and E-cadherin feedback. bioRxiv:2021.2002.2026.433093.

Chu, C.W., G. Masak, J. Yang, and L.A. Davidson. 2020. From biomechanics to mechanobiology: Xenopus provides direct access to the physical principles that shape the embryo. Curr Opin Genet Dev. 63:71–77.

Devitt, C.C., C. Lee, R.M. Cox, O. Papoulas, J. Alvarado, S. Shekhar, E.M. Marcotte, and J.B. Wallingford. 2021. Twinfilin1 controls lamellipodial protrusive activity and actin turnover during vertebrate gastrulation. Journal of Cell Science.

Fagotto, F., N. Rohani, A.S. Touret, and R. Li. 2013. A molecular base for cell sorting at embryonic boundaries: contact inhibition of cadherin adhesion by ephrin/ Eph-dependent contractility. Dev Cell. 27:72–87.

Fang, X., H. Ji, S.-W. Kim, J.-I. Park, T.G. Vaught, P.Z. Anastasiadis, M. Ciesiolka, and P.D. McCrea. 2004. Vertebrate development requires ARVCF and p120 catenins and their interplay with RhoA and Rac. The Journal of Cell Biology. 165:87–98.

Fernandez-Gonzalez, R., M. Simoes Sde, J.C. Roper, S. Eaton, and J.A. Zallen. 2009. Myosin II dynamics are regulated by tension in intercalating cells. Dev Cell. 17:736–743.

Finegan, T.M., N. Hervieux, A. Nestor-Bergmann, A.G. Fletcher, G.B. Blanchard, and B. Sanson. 2019. The tricellular vertex-specific adhesion molecule Sidekick facilitates polarised cell intercalation during Drosophila axis extension. PLOS Biology. 17:e3000522.

Fletcher, A.G., F. Cooper, and R.E. Baker. 2017. Mechanocellular models of epithelial morphogenesis. Philos Trans R Soc Lond B Biol Sci. 372.

Huebner, R.J., A.N. Malmi-Kakkada, S. Sarikaya, S. Weng, D. Thirumalai, and J.B. Wallingford. 2021a. Mechanical heterogeneity along single cell-cell junctions is driven by lateral clustering of cadherins during vertebrate axis elongation. eLife. 10:e65390.

Huebner, R.J., and J.B. Wallingford. 2018. Coming to Consensus: A Unifying Model Emerges for Convergent Extension. Dev Cell. 46:389–396.

Huebner, R.J., S. Weng, C. Lee, S. Sarikaya, O. Papoulas, R.M. Cox, E.M. Marcotte, and J.B. Wallingford. 2021b. Cell adhesions link subcellular actomyosin dynamics to tissue scale force production during vertebrate convergent extension. bioRxiv.

Irvine, K.D., and E. Wieschaus. 1994. Cell intercalation during Drosophila germband extension and its regulation by pair-rule segmentation genes. Development. 120:827–841.

Keller, R. 2002. Shaping the vertebrate body plan by polarized embryonic cell movements. Science. 298:1950–1954.

Keller, R., L.A. Davidson, and D.R. Shook. 2003. How we are shaped: the biomechanics of gastrulation. Differentiation. 71:171–205.

Keller, R., and J. Hardin. 1987. Cell behaviour during active cell rearrangement: evidence and speculations. J Cell Sci Suppl. 8:369–393.

Keller, R., J. Shih, and A. Sater. 1992. The cellular basis of the convergence and extension of the Xenopus neural plate. Dev Dyn. 193:199–217.

Keller, R., and A. Sutherland. 2020. Chapter Ten - Convergent extension in the amphibian, Xenopus laevis. In Current Topics in Developmental Biology. Vol. 136. L. Solnica-Krezel, editor. Academic Press. 271–317.

Keller, R., and P. Tibbetts. 1989. Mediolateral cell intercalation in the dorsal, axial mesoderm of Xenopus laevis. Dev Biol. 131:539–549.

Kim, H.Y., and L.A. Davidson. 2011. Punctuated actin contractions during convergent extension and their permissive regulation by the non-canonical Wnt-signaling pathway. J Cell Sci. 124:635–646.

Lecuit, T., and A.S. Yap. 2015. E-cadherin junctions as active mechanical integrators in tissue dynamics. Nature Cell Biology. 17:533–539.

Lienkamp, S.S., K. Liu, C.M. Karner, T.J. Carroll, O. Ronneberger, J.B. Wallingford, and G. Walz. 2012. Vertebrate kidney tubules elongate using a planar cell polarity-dependent, rosette-based mechanism of convergent extension. Nat Genet. 44:1382–1387.

Martin, A.C., M. Gelbart, R. Fernandez-Gonzalez, M. Kaschube, and E.F. Wieschaus. 2010. Integration of contractile forces during tissue invagination. J Cell Biol. 188:735–749.

Merkel, M., and M.L. Manning. 2017. Using cell deformation and motion to predict forces and collective behavior in morphogenesis. Semin Cell Dev Biol. 67:161–169.

Miao, H., and J.T. Blankenship. 2020. The pulse of morphogenesis: actomyosin dynamics and regulation in epithelia. Development. 147.

Mitrossilis, D., J. Fouchard, A. Guiroy, N. Desprat, N. Rodriguez, B. Fabry, and A. Asnacios. 2009. Single-cell response to stiffness exhibits muscle-like behavior. Proceedings of the National Academy of Sciences. 106:18243–18248.

Nishimura, T., H. Honda, and M. Takeichi. 2012. Planar cell polarity links axes of spatial dynamics in neural-tube closure. Cell. 149:1084–1097.

Novikova, E.A., and C. Storm. 2013. Contractile fibers and catch-bond clusters: a biological force sensor? Biophys J. 105:1336–1345.

Paulson, A.F., E. Mooney, X. Fang, H. Ji, and P.D. McCrea. 2000. Xarvcf, Xenopus Member of the p120 Catenin Subfamily Associating with Cadherin Juxtamembrane Region *. Journal of Biological Chemistry. 275:30124–30131.

Pfister, K., D.R. Shook, C. Chang, R. Keller, and P. Skoglund. 2016. Molecular model for force production and transmission during vertebrate gastrulation. Development. 143:715–727.

Rakshit, S., Y. Zhang, K. Manibog, O. Shafraz, and S. Sivasankar. 2012. Ideal, catch, and slip bonds in cadherin adhesion. Proceedings of the National Academy of Sciences. 109:18815.

Shih, J., and R. Keller. 1992. Cell motility driving mediolateral intercalation in explants of Xenopus laevis. Development. 116:901–914.

Shindo, A. 2018. Models of convergent extension during morphogenesis. Wiley Interdisciplinary Reviews: Developmental Biology. 7:e293.

Shindo, A., Y. Inoue, M. Kinoshita, and J.B. Wallingford. 2019. PCP-dependent transcellular regulation of actomyosin oscillation facilitates convergent extension of vertebrate tissue. Developmental Biology. 446:159–167.

Shindo, A., and J.B. Wallingford. 2014. PCP and septins compartmentalize cortical actomyosin to direct collective cell movement. Science. 343:649–652.

Sun, Z., C. Amourda, M. Shagirov, Y. Hara, T.E. Saunders, and Y. Toyama. 2017. Basolateral protrusion and apical contraction cooperatively drive Drosophila germ-band extension. Nat Cell Biol. 19:375–383.

Tada, M., and C.P. Heisenberg. 2012. Convergent extension: using collective cell migration and cell intercalation to shape embryos. Development. 139:3897–3904.

Tahinci, E., and K. Symes. 2003. Distinct functions of Rho and Rac are required for convergent extension during Xenopus gastrulation. Developmental Biology. 259:318–335.

Tamulonis, C., M. Postma, H.Q. Marlow, C.R. Magie, J. de Jong, and J. Kaandorp. 2011. A cell-based model of Nematostella vectensis gastrulation including bottle cell formation, invagination and zippering. Developmental Biology. 351:217–228.

Uyeda, T.Q.P., Y. Iwadate, N. Umeki, A. Nagasaki, and S. Yumura. 2011. Stretching Actin Filaments within Cells Enhances their Affinity for the Myosin II Motor Domain. PLOS ONE. 6:e26200.

Vanderleest, T.E., C.M. Smits, Y. Xie, C.E. Jewett, J.T. Blankenship, and D. Loerke. 2018. Vertex sliding drives intercalation by radial coupling of adhesion and actomyosin networks during Drosophila germband extension. eLife. 7:e34586.

Walck-Shannon, E., and J. Hardin. 2014. Cell intercalation from top to bottom. Nat Rev Mol Cell Biol. 15:34–48.

Wallingford, J.B., L.A. Niswander, G.M. Shaw, and R.H. Finnell. 2013. The continuing challenge of understanding, preventing, and treating neural tube defects. Science. 339:1222002.

Williams, M., W. Yen, X. Lu, and A. Sutherland. 2014. Distinct apical and basolateral mechanisms drive planar cell polarity-dependent convergent extension of the mouse neural plate. Dev Cell. 29:34–46.

Wilson, P.A., G. Oster, and R. Keller. 1989. Cell rearrangement and segmentation in Xenopus: direct observation of cultured explants. Development. 105:155–166.

Zaidel-Bar, R., G. Zhenhuan, and C. Luxenburg. 2015. The contractome – a systems view of actomyosin contractility in non-muscle cells. Journal of Cell Science. 128:2209–2217.

Zallen, J.A., and E. Wieschaus. 2004. Patterned gene expression directs bipolar planar polarity in Drosophila. Dev Cell. 6:343–355.

